# A hyperparameter benchmark of VAE-based methods for scRNA-seq batch integration

**DOI:** 10.64898/2026.02.10.705093

**Authors:** Mohamad Kassab, Luiz Maniero, Eduardo da Veiga Beltrame

## Abstract

We present the first systematic benchmark of model-architecture hyperparameters for variational autoencoder (VAE) methods for single-cell RNA-seq batch integration within scvi-tools, comparing scVI, MrVI, and LDVAE across four heterogeneous datasets under two feature regimes (all genes vs highly variable genes (HVGs)). We investigated 960 trainings (120 configurations) varying latent size and network depth/width, and evaluated with a standardized scIB metric suite covering batch removal and biological conservation (Batch ASW, PCR-batch, iLISI, graph connectivity, NMI, ARI, label ASW, isolated-label F1/ASW, cLISI, trajectory conservation), plus qualitative UMAP/t-SNE and PCA, random projection, and unintegrated baselines. Results show dataset-dependent trade-offs: scVI performs best overall via stronger batch correction; LDVAE can better preserve biological structure in some datasets; MrVI is stable and excels at batch correction in multi-protocol settings but is more resource-intensive. HVG-only training generally outperforms full-gene training for all models. Hyperparameter analysis suggests moderate-to-high latent dimensionality (>30) often gives the best balance; sensitivity to latent size tracks dataset heterogeneity (tissues, labs, chemistries, gene coverage), with larger latents improving batch mixing but sometimes reducing biological conservation. We provide model- and dataset-specific guidelines for practical defaults and tuning of VAE-based integration in single-cell studies. Reproducibility code is available on GitHub at: https://github.com/Kassab11/A-hyperparameter-benchmark-of-VAE-based-methods-for-scRNA-seq-batch-integration

## Introduction

Single-cell gene expression resources have expanded rapidly, enabled by high-throughput methods that profile thousands to millions of cells per experiment [2–8]. This scale is visible in public portals: by October 2024, CELLxGENE reported 169.3 million cells, including 93.6 million unique cells across >2,000 cell types [9], and the Human Cell Atlas portal listed 64.4 million cells by September 2025 [10]. Tahoe-100M further exemplifies this growth with 100 million perturbation-resolved profiles [11]. However, the diversity of scRNA-seq chemistries and platforms—varying capture efficiency, read coverage, throughput, and workflows—introduces batch effects that can obscure biological signal [12–14] and drive clustering by run, chemistry, or laboratory without careful design and integration [15].

Batch-effect correction methods have been extensively studied: many works introduce new algorithms and compare them to existing approaches [12, 16–23], others provide broad benchmarks [1, 15, 24–28] or evaluation metrics [29, 30]. Yet, to our knowledge, no prior work has systematically benchmarked architecture hyperparameter configurations of variational-autoencoder (VAE)–based methods specifically for batch correction.

Within scvi-tools, VAE models remain among the most widely used probabilistic approaches for scRNA-seq integration and analysis, combining strong performance with modest compute and mature tooling [31–39]. Transformer-based single-cell foundation models (FMs) have also emerged, e.g., scFoundation [40] and AIDO.Cell [41]. AIDO.Cell was pre-trained on 50M cells and required large-scale training (256 H100 GPUs over three days for the 100M-parameter FM, and eight days for the 650M-parameter FM) [41]. However, independent zero-shot evaluations report that popular single-cell FMs can underperform simpler approaches, including scVI, on PBMC68k identification tasks by NMI/ARI [41], keeping scVI and its variants (MrVI for cohort-level effects; LDVAE for interpretability via a linear decoder) practical choices. See Methods for more details on the selected models in this study.

Furthermore, many studies that utilize scVI in their analysis tend to use the default settings (latent dimensionality of 10, hidden layer of 1, and 128 nodes per layer) [35–39, 42]. Some studies report using different parameters [31, 33, 34]; however, they tend to use a single combination and do not provide a clear reasoning for the choice of parameters.

To address this gap, we benchmarked scVI, MrVI, and LDVAE for scRNA-seq integration, spanning 120 architecture configurations (30 scVI, 60 MrVI, 30 LDVAE) under two gene-selection regimes (full genes vs top 5,000 HVGs) across four datasets, totaling 960 training runs. Our benchmarking pipeline is described in Figure 1. We varied four core hyperparameters as shown in Table 1 to characterize operating regimes and sensitivity to latent size, depth, width, and (for MrVI) the sample-aware latent—choices that directly govern the batch-versus-biology trade-off. We do not claim that optimization parameters (learning rate, batch size, dropout) are not important; rather, they are supported by established guidance [43–48] and are outside of our scope. Integration quality was assessed with Batch ASW, PCR batch, iL-ISI, graph connectivity, NMI, ARI, label ASW, isolated label F1, isolated label ASW, cLISI, and trajectory conservation. Isolated label F1, isolated label ASW, and trajectory conservation were unavailable for some datasets due to label/trajectory availability. We additionally used t-SNE and UMAP for qualitative assessment and compared against unintegrated, random projection, and PCA baselines. Datasets included the Human Immune dataset from OpenProblems [1], Zenodo 8020792 (24 PBMC Samples) [49], Zenodo 11100300 (18 PBMC Samples) [50], and Tabula Muris [51], See Datasets for more details on the datasets used in this benchmarking study.

**Table 1.** Summary of hyperparameter configurations explored for each model. ✓indicates applicability.

| Hyperparameter | Values | scVI | MrVI | LDVAE |
| --- | --- | --- | --- | --- |
| $n_{\text{latent}}^Z$ | 10, 20, 30, 40, 50 | ✓ | ✓ | ✓ |
| $n_{\text{latent}}^U$ | 10, 20 | | ✓ | |
| $n_{\text{layers}}$ | 1, 2, 3 | ✓ | ✓ | ✓ |
| $n_{\text{hidden}}$ | 128, 256 | ✓ | ✓ | ✓ |

**Table 2.**
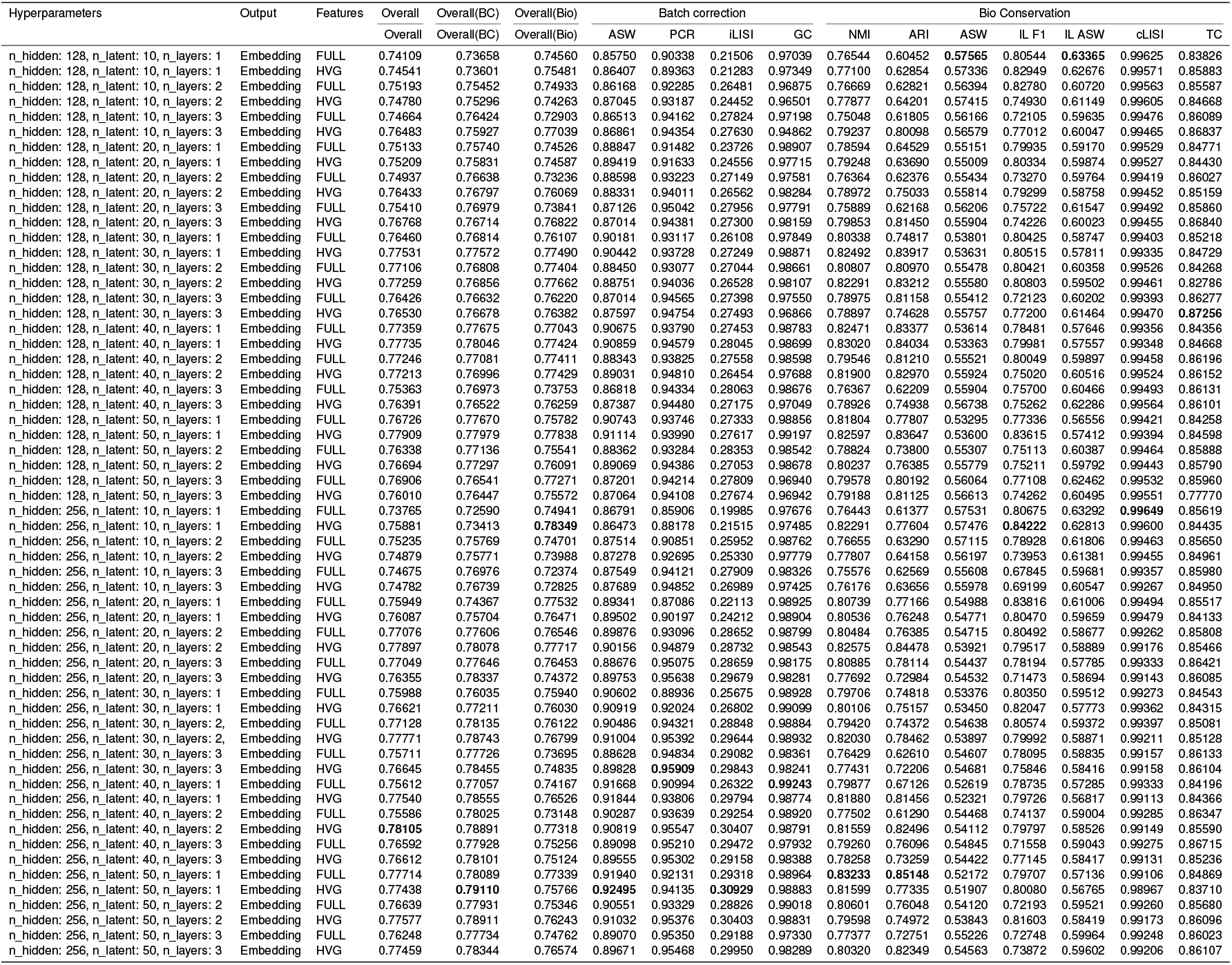
Performance of scVI on the Human Immune Dataset.

**Figure 1.**
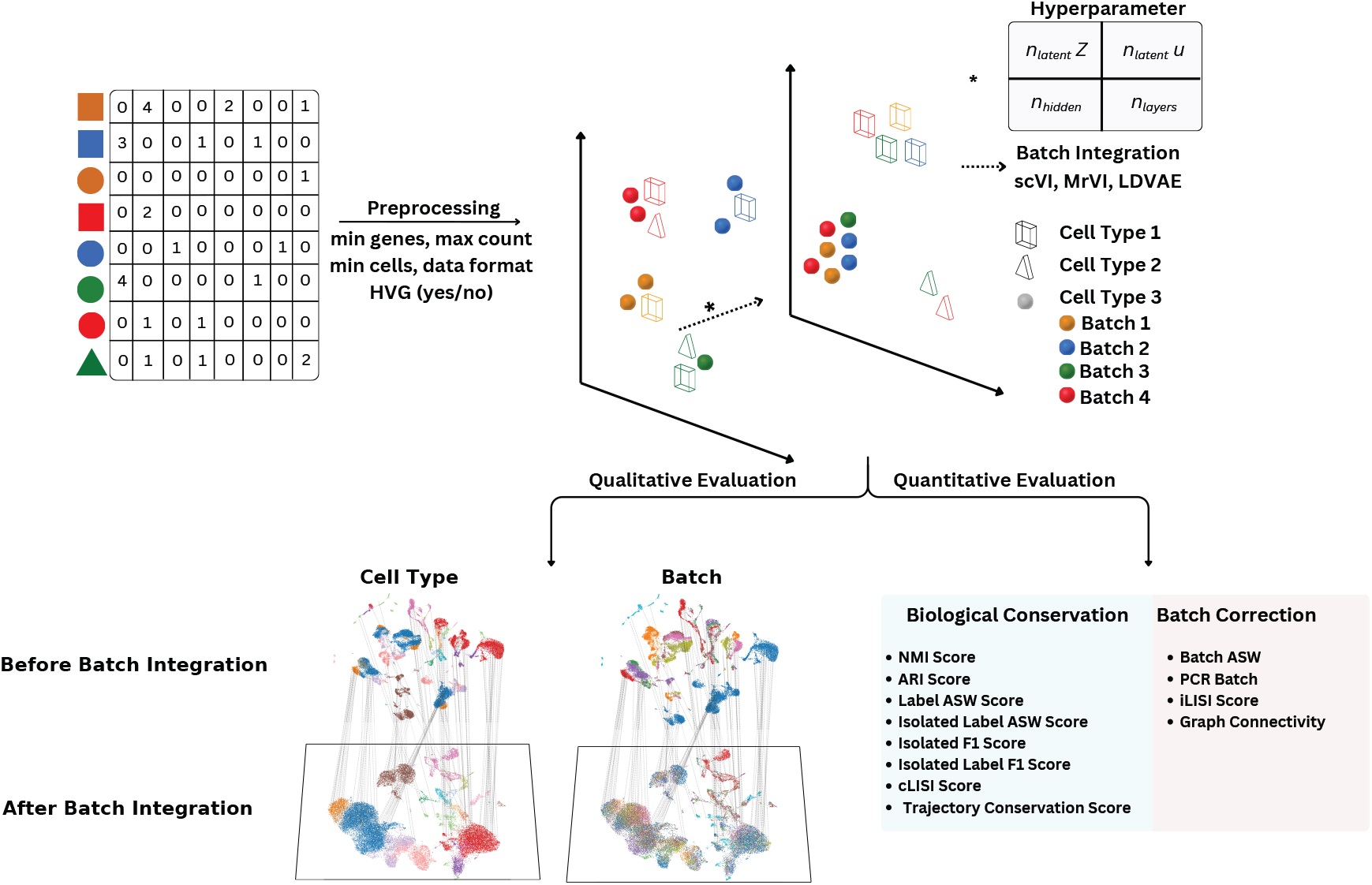
Hyperparameter benchmarking workflow for VAE-based scRNA-seq data integration methods. (i) Starting from a gene-by-cell count matrix covering multiple experimental batches (colours) and annotated cell types (shapes), data were preprocessed by filtering (minimum genes per cell, maximum counts per cell, minimum cells per gene). Analyses were run under two gene-selection regimes: full gene sets or the top 5,000 HVGs. (ii) VAE-based batch-integration models (scVI, MrVI and LDVAE) were trained across a grid of architectural hyperparameters to probe operating regimes and the batch–biology trade-off: shared latent dimensionality (nlatent, Z), MrVI sample-aware latent dimensionality (nlatent, U), hidden-layer width (nhidden) and network depth (nlayers). Training yields a low-dimensional embedding in which cells of the same type are aligned across batches while preserving biological separation (dashed arrow). (iii) Qualitative evaluation was performed by visualizing embeddings (UMAP and t-SNE) coloured by cell type or batch before and after integration. (iv) Quantitative evaluation, through scIB metrics [1], captured both biological conservation and batch correction, using normalised mutual information (NMI), adjusted Rand index (ARI), label average silhouette width (label ASW), isolated label ASW, isolated label F1, cell-type local inverse Simpson’s index (cLISI) and trajectory conservation (when available), alongside batch ASW, principal component regression on batch (PCR batch), integration LISI (iLISI) and graph connectivity.

## Results

The following section reports results for each dataset across the three models, condensed for readers seeking practical tuning guidelines for the studied VAEs. All metrics follow the definitions in [1]. We compute an overall score as the mean of two group-wise averages: (i) biological conservation and (ii) batch correction. To reflect the primacy of preserving biological signal, we aggregate seven biological metrics and four batch-correction metrics, since removing batch noise without maintaining biology is not a desirable outcome.

### Immune Datasets

Figure 2 compares one–parameter sweeps in which two hyperparameters are fixed while the third is varied; a detailed paired, cross–hyperparameter analysis along with qualitative analysis is provided in Human Immune Datasets - Detailed Results. Visually, a moderate to high latent dimensionality *n*_latent*Z*_ usually strikes the best balance for scVI, MrVI, and LDVAE. Sensitivity to *n*_latent*Z*_ is strongest in the Human Immune dataset: pushing *n*_latent*Z*_ to the high end boosts the batch composite (BC) but tends to soften the biology composite (Bio) for MrVI and LD-VAE. Increasing the hidden width *n*_hidden_ yields at most small, marginal gains in a few settings and is otherwise flat or mildly negative. Increasing depth *n*_layers_ often lifts BC while generally reducing overall (Bio) at higher *n*_layers_, producing a clear trade–off; among the models, LDVAE appears least sensitive to this depth change.

**Figure 2.**
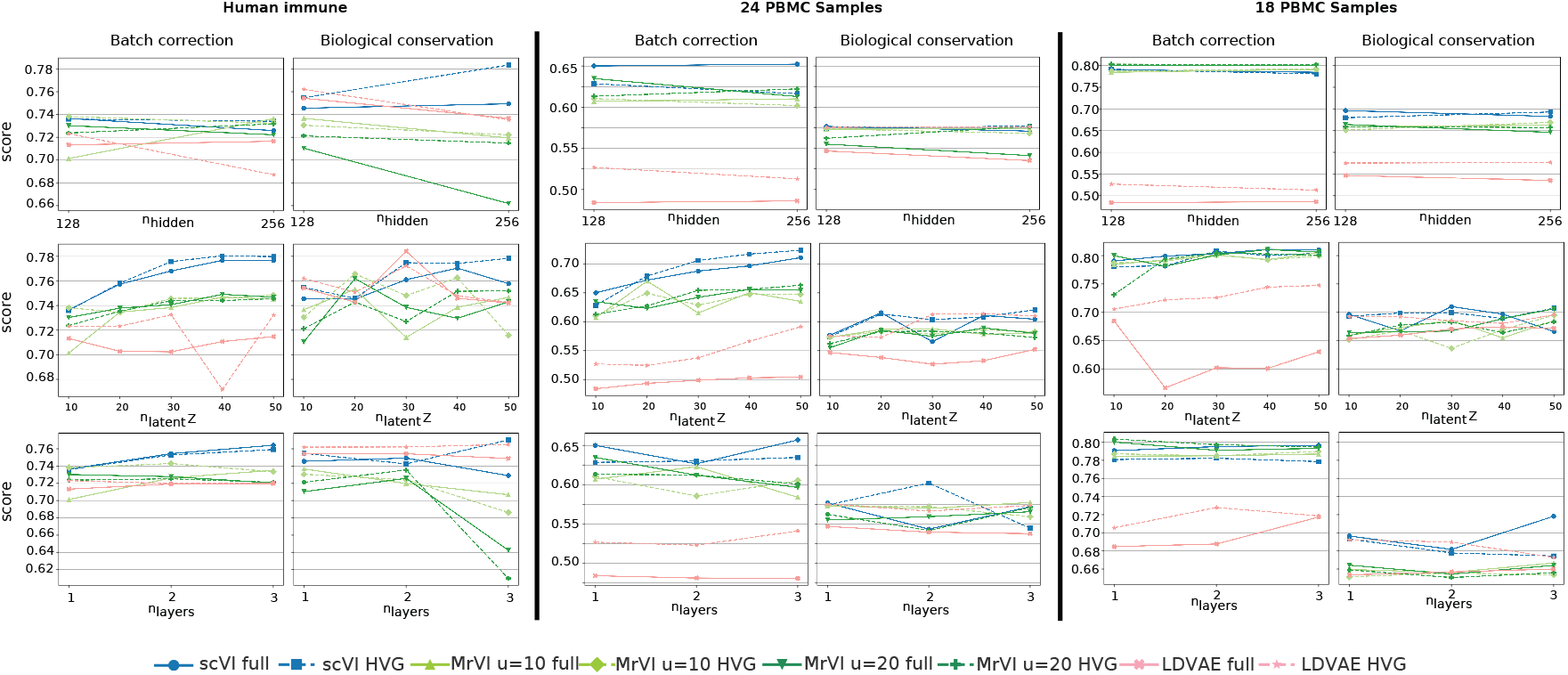
Hyperparameter sensitivity of scVI, MrVI and LDVAE across different datasets. Columns show results for three datasets (left to right): Human Immune (OpenProblems), Zenodo 8020792 (24 PBMC samples) and Zenodo 11100300 (18 PBMC samples). For each dataset, the left subpanels report the overall batch-correction score and the right subpanels report the overall biological-conservation score (higher is better for both). Rows show one-dimensional sweeps through the architecture grid in which two architectural hyperparameters are held fixed while the third is varied: hidden-layer width (nhidden; top), shared latent dimensionality (nlatent,Z; middle) and network depth (nlayers; bottom). MrVI is shown for two settings of the sample-aware latent (u = nlatent,U = 10 or 20). Solid lines denote models trained using the full gene set, and dashed lines denote models trained using the top 5,000 HVGs; markers/colours indicate model family as in the key. Overall batch-correction scores aggregate batch-focused metrics, whereas overall biological-conservation scores aggregate biology-focused metrics.

Figure 3 shows the best-performing configuration of each model across the immune datasets, alongside Unintegrated, PCA, and Random Projection baselines. Overall, scVI consistently leads: it achieves the strongest batch correction on Human Immune and Zenodo 8020792 (24 PBMC Samples) (with HVG), and on Zenodo 11100300 (18 PBMC Samples) it delivers top biological conservation with a batch score close to the best. MrVI is notably steady across datasets for both Full and HVG features, with batch and biology scores that are closely matched. LDVAE exhibits data set-specific gains, most prominently in Human Immune, where it attains the highest biology score, and benefits highly from the HVG features in the remaining datasets.

**Figure 3.**
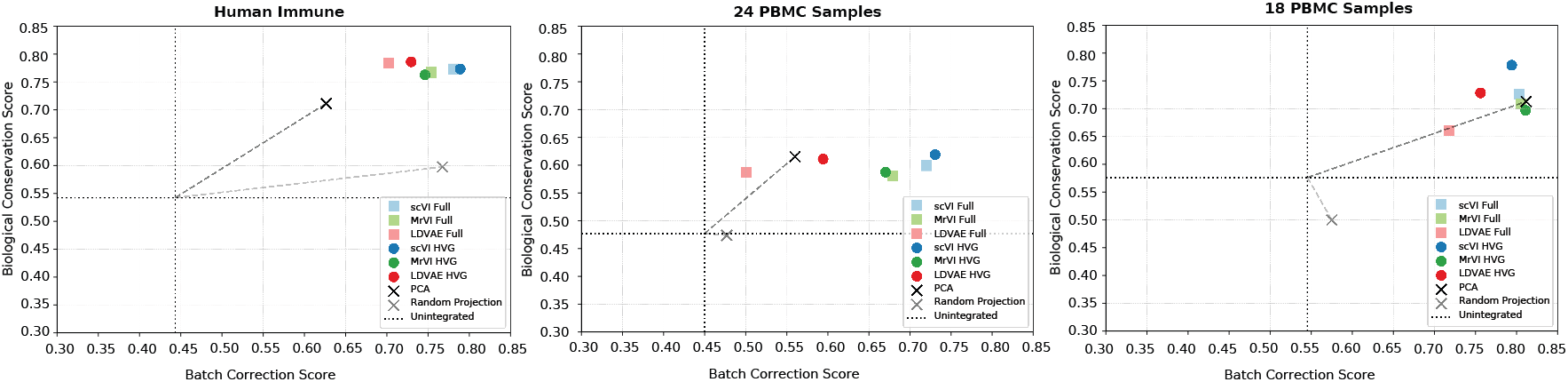
Best-achieved batch correction versus biological conservation across VAE-based models. Scatter plots of the overall batch-correction score (x axis) against the overall biological-conservation score (y axis) for the Human Immune (OpenProblems), 24 PBMC Samples (Zenodo 8020792); and 18 PBMC Samples (Zenodo 11100300). Coloured points denote the best-performing configuration for each integration model (scVI, MrVI and LDVAE). Squares indicate models trained on the full gene set, and circles indicate models trained on the top 5,000 HVGs. Crosses show non-integration baselines (PCA and a random projection of the same input). Dotted vertical and horizontal lines mark the corresponding unintegrated baseline scores, and dashed connectors indicate the shift from the unintegrated baseline to PCA or random projection. Points closer to the upper-right corner indicate stronger batch mixing while preserving biological structure.

### Tabula Muris Dataset

We use this dataset to validate our findings and assess whether the same hyperparameter effects hold. As shown in Figure 4, the first panel demonstrates that sweeping one hyperparameter while holding the other two fixed leads to the same conclusion as before: increasing the latent dimension *z* beyond 30 improves both batch correction and biological conservation. The second panel further indicates that scVI attains the highest overall score. Notably, because Tabula Muris is highly heterogeneous and involves two protocols and 154 cell types (51 immune and 103 non-immune), the sample-aware covariate is influential; accordingly, MrVI excels in batch correction, which is consistent with its design.

**Figure 4.**
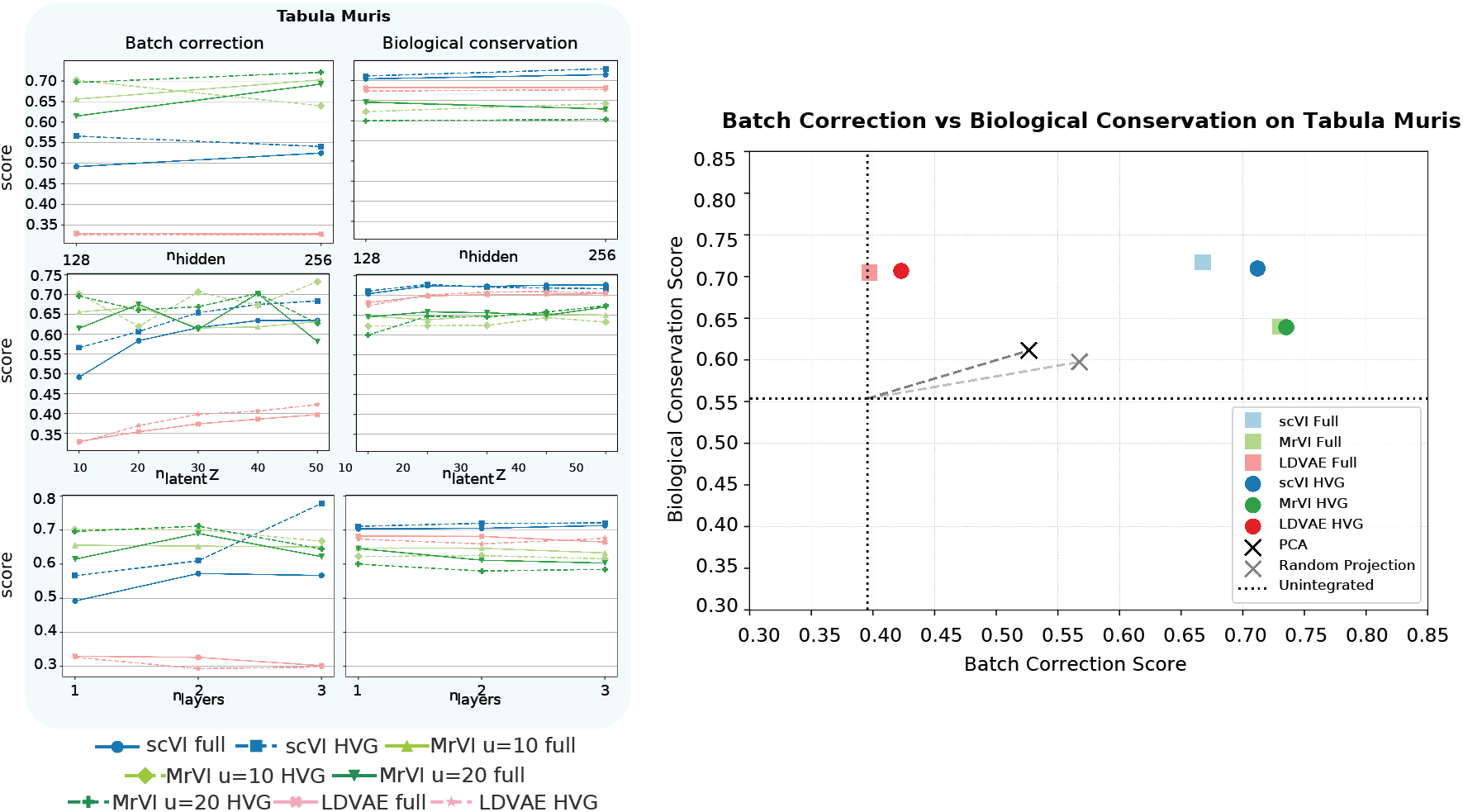
Hyperparameter sensitivity and batch-biology trade-off on Tabula Muris dataset. (i) Left subpanels report the overall batch-correction score and the right subpanels report the overall biological-conservation score (higher is better for both). Rows show one-dimensional sweeps through the architecture grid in which two architectural hyperparameters are held fixed while the third is varied: hidden-layer width (nhidden; top), shared latent dimensionality (nlatent,Z; middle) and network depth (nlayers; bottom). MrVI is shown for two settings of the sample-aware latent (u = nlatent,U = 10 or 20). Solid lines denote models trained using the full gene set, and dashed lines denote models trained using the top 5,000 HVGs; markers/colours indicate model family as in the key. Overall batch-correction scores aggregate batch-focused metrics, whereas overall biological-conservation scores aggregate biology-focused metrics. (ii) Overall batch-correction versus biological-conservation scores on Tabula Muris dataset for the best-performing configuration of each model under each gene-selection regime. Squares indicate full-gene models and circles indicate HVG models. Crosses denote non-integration baselines (PCA and a random projection of the same input features). Dotted vertical and horizontal lines mark the unintegrated baseline scores, and dashed connectors indicate the corresponding shifts from the unintegrated baseline to PCA or random projection. Points closer to the upper-right indicate stronger batch mixing while better preserving biological structure.

**Figure 5.**
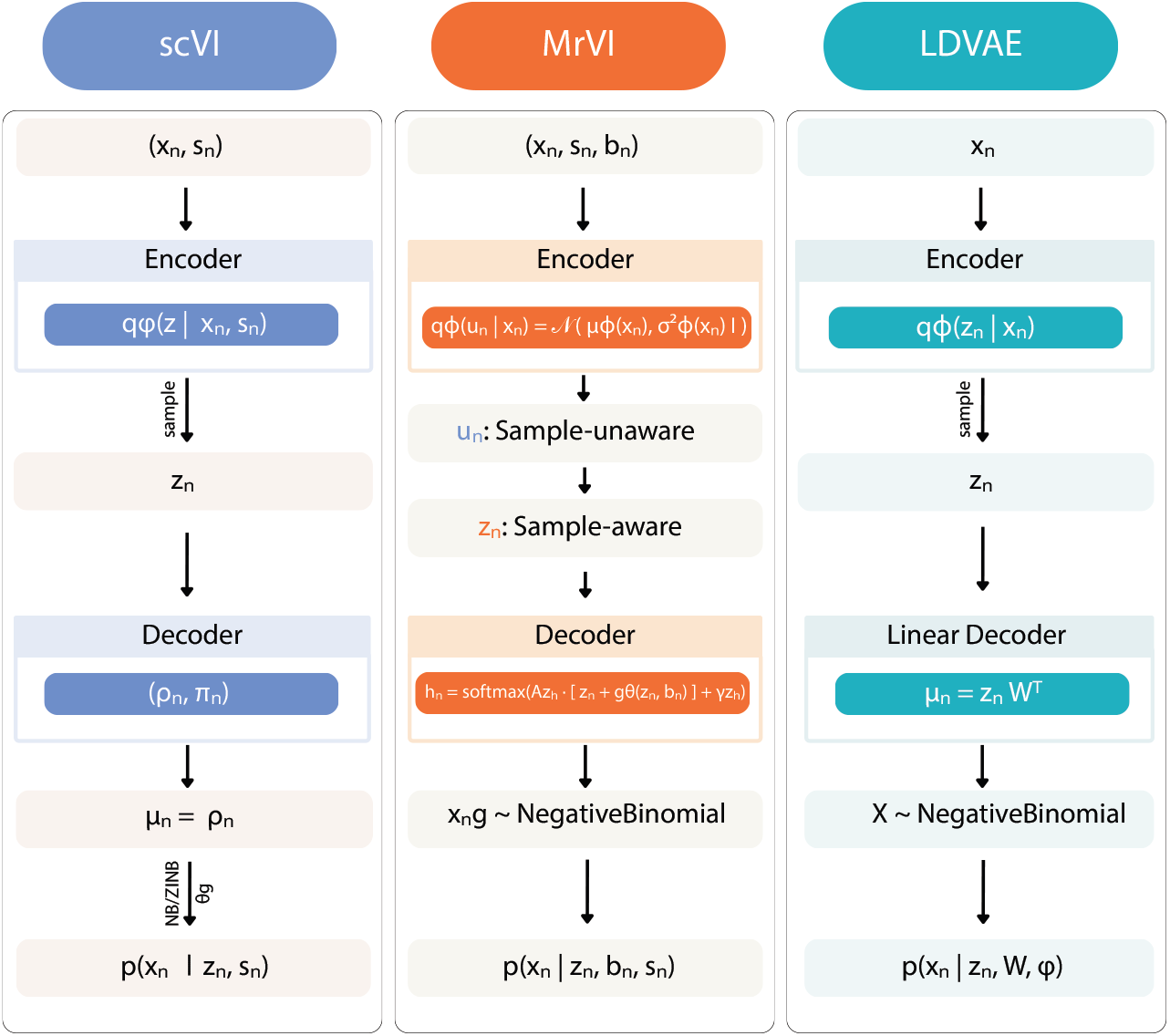
Graphical representations of probabilistic models scVI, MrVI, and LDVAE [53–55]. Each panel shows the flow from observed single-cell RNA-seq counts to latent representations (via an encoder) and back to a count likelihood (via a decoder). **scVI:** x_n_are observed gene counts for cell n and s_n_ are observed covariates (typically batch). The encoder approximates posteriors over a latent cell state z_n_. The decoder outputs gene-expression proportions ρ _n_and (optionally) a zero-inflation parameter π.The mean count for gene g in cell n is µ _ng_= ρ _ng_, and θ _g_denotes gene-specific dispersion used in the NB/ZINB likelihood for x_ng_. **MrVI:** x_n_ are observed gene counts,s_n_ is the sample ID, and b_n_ are nuisance covariates (e.g., batch). The encoder defines | x_n_) 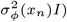 where u_n_ is a sample-unaware latent representation, µ_φ_(x_n_) and 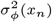 are encoder-predicted mean and variance, and I is the identity matrix (diagonal covariance). A sample-aware latent representation z_n_ is formed using s_n_. The decoder produces gene-expression proportions h_n_ via a linear map A_zh_, a nuisance correction term g_θ_(z_n_, b_n_), and bias γ _zh_; observed counts x_ng_ are modeled with a Negative Binomial likelihood. **LDVAE:** x_n_ are observed gene counts for cell n. The encoder posterior q_φ_(z_n_ x_n_) (parameters φ) yields a low-dimensional cell representation z_n_. The linear, interpretable decoder outputs µ_n_ = z_n_W ^⊤^, where W is the gene-loading matrix.

## Discussion

Our analysis shows that the most compute-efficient way to add model capacity is to increase the latent dimension z, as it tends to improve performance with the smallest hit to training time and memory (especially for scVI). Whereas increasing hidden width, depth, batch size, or the number of genes is typically much more expensive. A detailed discussion of computational complexity is provided in Computational Complexity. We further analyze score-complexity trade-offs in Score-Complexity Trade-offs.

### scVI

Among the evaluated models, scVI is the least expensive to tune in terms of the latent dimension z. Across four datasets spanning a wide range of cell types and technologies, scVI consistently outperformed the alternatives, making it a strong default choice for scRNA-seq analysis. In practice, we recommend tuning the latent dimension *z* first—using highly variable genes (HVGs)—as this offers the best accuracy–efficiency trade-off and minimizes computational cost. Across all datasets we observe that increasing the latent dimensionality *z* consistently improves both batch integration and biological conservation; in practice, we recommend moderate–to–high *z* (30 - 50). In contrast, increasing the hidden dimension from *H* = 128 to *H* = 256 yields only marginal gains, especially once *z* > 30 (see Fig. 8) and can even reduce biological conservation, while adding depth typically provides small benefits at best and more often degrades performance at higher layer counts. It is important to note that scVI comprises an MLP encoder with two output heads and one decoder, so the operations that scale with *z* are confined to *H*→ *z* (two heads) and *z* → *H*, giving train-time complexity 𝒪 (*BHz*), which is linear in *z*. In contrast, the genes facing layers *G* → *H* and *H* → *G* scale as 𝒪 (*BGH*), and each additional hidden layer contributes 𝒪 (*BH*^2^). Since typical regimes have *z* in the tens, *H* in the hundreds, and *G* in the thousands, the marginal compute of expanding *z* is negligible, making *n*_latent_ a compute efficient lever that often improves accuracy without meaningful run-time or memory penalties.

### MrVI

We observe that setting the latent *z* dimensionality in the moderate–high range yields the best results; increasing *n*_layers_ offers little benefit and, in the Human Immune dataset, setting *n*_layers_ = 3 steeply decreases overall biological conservation. The t-SNE maps (Fig.) show improvement when increasing the auxiliary latent *u* from 10 to 20, and in certain settings at higher *n*_latent_*z, u* = 20 preserves the compactness of clustering. We therefore recommend, for MrVI with HVG features, using moderate–high *n*_latent_ ∈ [20, 40] with *u* = 20, a shallow depth *n*_layers_ = 1, and *n*_hidden_ = 128. Furthermore, from the Tabula Muris dataset, we see that MrVI exhibited strong stability and excelled at batch correction on highly heterogeneous datasets (154 cell types), largely due to its architecture, which includes a sample-aware covariate capable of accommodating multiple protocols. We therefore recommend MrVI for complex, multi-protocol datasets. However, it is important to note that MrVI is the most computationally expensive model.

### LDVAE

Excels at interpretability and scaling because it allows to visualize how latent dimensions link to genes, but has less capacity to model complex, nonlinear manifolds and to integrate datasets with complex batch effects. It performs best when heterogeneity is dominated by broad lineages (as in Human Immune), where gene programs are nearly axis-aligned and a linear decoder cleanly preserves biology across batches. By contrast, on single-tissue PBMC cohorts rich in donor/time effects and fine subtype boundaries and on Tabula Muris, which is a multi-tissue, cross-protocol atlas, nonlinearity buys accuracy, and LD-VAE’s linearity becomes a bottleneck.

Our results indicate that HVG is a preferable gene-selection strategy, consistent with prior studies [1, 52], thereby serving as a sanity check for our analysis. In general, we recommend scVI as the go-to model across datasets; although many studies utilize scVI in their analysis, they tend to use the default settings (*n*_latent_=10, *n*_layers_=1, *n*_hidden_=128), as cited in the introduction, which is not the best performing choice, and this reliance on defaults overlooks the fact that parameter tuning can yield improved results, as our analysis demonstrates. A strong, compute-efficient starting point is to use HVG features and a moderate to high latent size *z* in the range 30–50, a shallow encoder with *n*_layers_=1, 2, and *n*_hidden_=128.

## Conclusion

In this study, we systematically benchmarked VAE-based batch-integration methods (scVI, MrVI, LDVAE) across three datasets by sweeping feature sets (HVG vs. Full), latent size, network depth, and width. We found that widely used defaults are not optimal: HVG features and a moderate to high latent dimension (30 to 50) improve the overall quality of integration. scVI emerges as the most reliable general choice, LDVAE best preserves biology in the most heterogeneous dataset, and MrVI benefits from *u*=20 and larger *z* but adds computational overhead. Practically, we recommend adopting HVG by default, first tuning *z*, keeping depth shallow and only widening when necessary, yielding a robust and compute-efficient recipe for single-cell integration.

## Methods

Figure shows the graphical models used in this study, including scVI [53], LDVAE [55], and MrVI [54]. All of these models are considered as deep-generative models for scRNA-seq data, but differ in focus and interpretability. scVI (single cell variational inference) is a hierarchical Bayesian VAE that models gene counts with a zero-inflated negative binomial distribution, accounts for batch effects, and learns a low-dimensional latent space for tasks such as normalization, clustering, and differential expression [53]. LDVAE (Linearly Decoded VAE) is built directly on scVI, but replaces the non-linear decoder with a linear factor model, sacrificing reconstruction accuracy for interpretability, where each latent dimension corresponds to a gene program that can be biologically interpreted as coexpression axes [55]. MrVI (Multi-resolution Variational Inference) generalizes scVI to cohort-level studies by introducing hierarchical latent variables where one latent space captures cell states independent of samples, and another that incorporates sample-level covariates [54].

### scVI [53]

#### Observed data and annotations

For cells *n* = 1, …, *N* and genes *g* = 1, …, *G*, let *x*_*ng*_∈ℕbe observed counts. Each cell has a (known) batch annotation *s*_*n*_ ∈ {1,…, *B*}. The model conditions on *s*_*n*_.

#### Latent variables

scVI introduces a low-dimensional biological latent *z*_*n*_ ∈ℝ^*d*^ and a cell-specific library-size factor *l*_*n*_ > 0 (on the log scale). Priors:

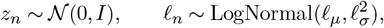

with batch-specific hyperparameters 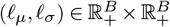

### Decoder parameterization (batch-conditioned)

Given (*z*_*n*_, *s*_*n*_), scVI decodes gene-wise parameters via neural networks:

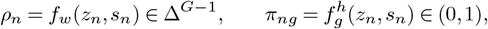

where *ρ*_*n*_ is a batch-corrected, normalized vector of transcript proportions (summing to one) and *π*_*ng*_ is the zero-inflation probability. The mean of the NB component is *µ*_*ng*_ = *f*_*n*_ *ρ*_*ng*_ with gene-specific inverse-dispersion *θ*_*g*_ > 0.

### Generative process

For each pair (*n, g*), the model draws

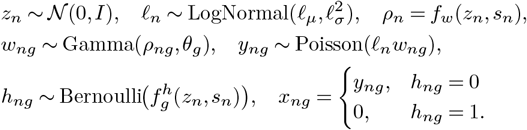

Marginalizing (*w*_*ng*_, *y*_*ng*_) yields a zero-inflated negative binomial (ZINB) likelihood for *x*_*ng*_ with mean *µ*_*ng*_ = *f*_*n*_*ρ*_*ng*_,dispersion *θ*_*g*_, and zero-inflation *π*_*ng*_.

### Approximate posterior (amortized VI)

Encoders parameterize a mean-field variational family

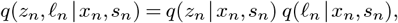

with *q*(*z*_*n*_|·) Gaussian and *q*(*f*_*n*_|·) log-normal. The evidence lower bound is

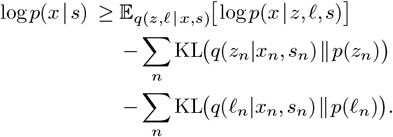

with {*θ*_*g*_*}* optimized as global variables. Training uses reparameterization and mini-batching.

### How batch integration is realized in the model

Batch enters the generative path only through the decoder conditionals *f*_*w*_(·, *s*_*n*_) and *f*^*h*^(·, *s*_*n*_). This induces a latent *z*_*n*_ that emphasizes biology while the decoder accounts for batch-dependent shifts in the observation model.

### LDVAE [55]

#### Model description

LDVAE replaces scVI’s non-linear decoder with a *linear* reconstruction map, yielding logits 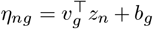 and proportions *h*_*n*_ = softmax(*η*_*n*_), so rows of *V* act as interpretable gene loadings directly linking latent axes to co-expressed “gene programs.” This design deliberately trades a modest increase in reconstruction error for interpretability in very large datasets. In the paper’s formulation, LDVAE is demonstrated in a single-batch setting with zero-inflation deactivated while retaining the scVI NB count model with a library-size factor.

### MrVI [54]

#### Observed variables and covariates

For cell *n*∈ *{*1, …, *N}* and genes *g {*1, …, *G}*, let counts be *x*_*ng*_ ∈ ℕ. Each cell has a *target* sample ID *s*_*n*_ ∈{1, …, *S*} and a *nuisance* covariate *b*_*n*_ ∈{1, …, *B}* (e.g. batch/chemistry).

#### Latent structure (two–level representation)

MrVI introduces (i) a *sample–unaware* latent 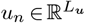 that captures cell state and is independent of (*s*_*n*_, *b*_*n*_), and (ii) a *sample–aware* latent 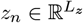 that augments *u*_*n*_ with sample effects while remaining independent of *b*_*n*_. Priors:

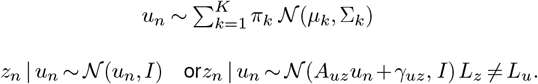

This hierarchy makes *u*_*n*_ the harmonized cell-state space and *z*_*n*_ the sample-conditioned refinement.

#### Decoder with nuisance conditioning and NB likelihood

Let *g*_*θ*_(*z*_*n*_, *b*_*n*_) denote a multi-head attention module that injects nuisance effects. Define logits for gene-wise normalized expression

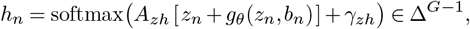

with learned 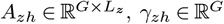 Given a size factor *l*_*n*_ > 0 (taken as the library size in the paper) and genewise inverse dispersion *r*_*ng*_ *≥*0, the observation model is Negative Binomial

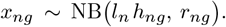

Thus nuisance *b*_*n*_ affects only the observation path via *g*_*θ*_, while sample effects enter through *z*_*n*_.

#### Joint distribution

Let 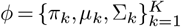 and *ψ* = {*A*_*uz*_, *γ*_*uz*_, *A*_*zh*_, *γ*_*zh*_, *θ, r*} denote parameters. The generative model factorizes as

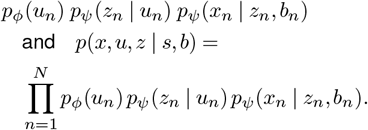

#### Variational inference and ELBO

With amortized posteriors *q*_*λ*_(*u*_*n*_| *x*_*n*_) and *q*_*λ*_(*z*_*n*_| *x*_*n*_, *s*_*n*_), the evidence lower bound is

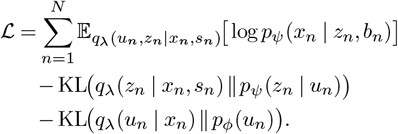

optimized over (*λ, φ, ψ*) with reparameterization gradients and mini-batching.

### Datasets

The choice of datasets in our study provides a rigorous benchmark for batch integration because they span complementary sources of variation. Together, they capture cross-study heterogeneity, concentrate many batches within a unified protocol, and incorporate repeated sampling over time. This combination challenges methods to handle differences driven by study design, run-and-handling effects, and temporal shifts. Each dataset includes explicit batch identifiers and harmonized annotations, allowing a consistent application of batch mixing and structure preservation metrics. The dataset sizes balance realism with tractability, enabling exhaustive hyper-parameter sweeps and repeated trials, and the shared file format with standardized metadata supports reproducible pipelines and fair comparisons. All used datasets underwent standardized preprocessing steps, including removal of low-quality cells (bottom 1% gene count, top 1% total count), high mitochondrial content cells, and duplicates. Genes expressed in fewer than three cells are also filtered. Filtering thresholds were defined per batch or library based on each dataset’s distribution.

### Human Immune dataset

The OpenProblems-scIB benchmark serves as a widely adopted reference for evaluating computational methods in the integration of data scRNA-seq. It aggregates immune cells from peripheral blood and bone marrow in five datasets and ten batches, generated using various versions of 10X Genomics (v2 and v3 chemistries) and Smart-seq2 technologies, as illustrated in Figure 12 in the Appendix. This dataset includes 33,506 cells and 12,303 genes, provided in AnnData format. After preprocessing, gene and total count thresholds varied across batches, generally ranging from 400–2700 genes and 8,000–890,000 total counts.

### Zenodo 11100300 (18 PBMC Samples)

A longitudinal scRNA-sequence dataset of peripheral blood mononuclear cells from 2 donors with myalgic encephalomyelitis/chronic fatigue syndrome, collected before, during, and after an antibiotic-induced remission event; it contains 55,260 cells and 36,601 genes across 4 batches, generated using 10x Genomics, and is provided in AnnData format as illustrated in Figure 13. After preprocessing, gene and total count thresholds ranged between 380–490 genes and 15,000–18,000 total counts. This dataset contains 18 PBMC samples.

### Zenodo 8020792 (24 PBMC Samples)

A capillary-blood PBMC scRNA-seq resource aggregating 28 samples from 3 donors across 14 experimental batches, generated using 10x Genomics v3.1 chemistry with MULTI-seq barcodes; it includes 76,535 cells and 36,601 genes and is provided in AnnData format as illustrated in Figure 14. This dataset includes a four-level hierarchical cell type annotation, we have used the cell type level 3 label to ensure a fine but stable granularity for biological conservation assessment. After preprocessing, thresholds ranged from 1,000–1,700 genes and 14,000–31,000 total counts, and were additionally filtered to retain only PBMCs based on cell type annotations. The final number of samples after preprocessing is 24 PBMC samples.

### Tabula Muris

The Tabula Muris atlas, a cross-tissue mouse singlecell RNA-seq resource profiling on the order of 10^5^ cells across roughly 20 organs using two complementary modalities, droplet-based 3^’^ UMI (10x) for breadth and FACS/Smart-seq2 for depth, yielding rich coverage of immune and non-immune compartments [51]. The dataset includes 356,213 cells and 20,116 genes, as illustrated in Figure. 15. The atlas includes 51 immune cell-type labels (e.g., T/B/NK, myeloid and dendritic subsets, tissue-resident macrophages) and 103 non-immune labels (epithelial, endothelial/vascular, stromal/mesenchymal, neural, muscle, hepatic, renal, pancreatic, etc.), for a total of about 154 distinct labels.

### Human Immune Datasets - Detailed Results

#### Qualitative Comparison

The t-SNE in figure 6, and the UMAP provided in Figure 7 compare the best performing configuration of scVI, MrVI, and LDVAE in Human Immune, Zenodo 8020792 (24 PBMC Samples), and Zenodo 11100300 (18 PBMC Samples) datasets, with each method colored by cell type and batch. Across datasets, scVI consistently produces the most compact within–type clusters with clear boundaries, while maintaining strong cross-batch overlap. MrVI clusters cells correctly by type most of the time, but we can still see traces of batch effect in the embedding, as shown in Zenodo 8020792 (24 PBMC Samples). LDVAE performs well on Human Immune, where clusters are recognizable and batches mix reasonably. However, in Zenodo 8020792 (24 PBMC Samples) it shows poor performance in clustering.

**Figure 6.**
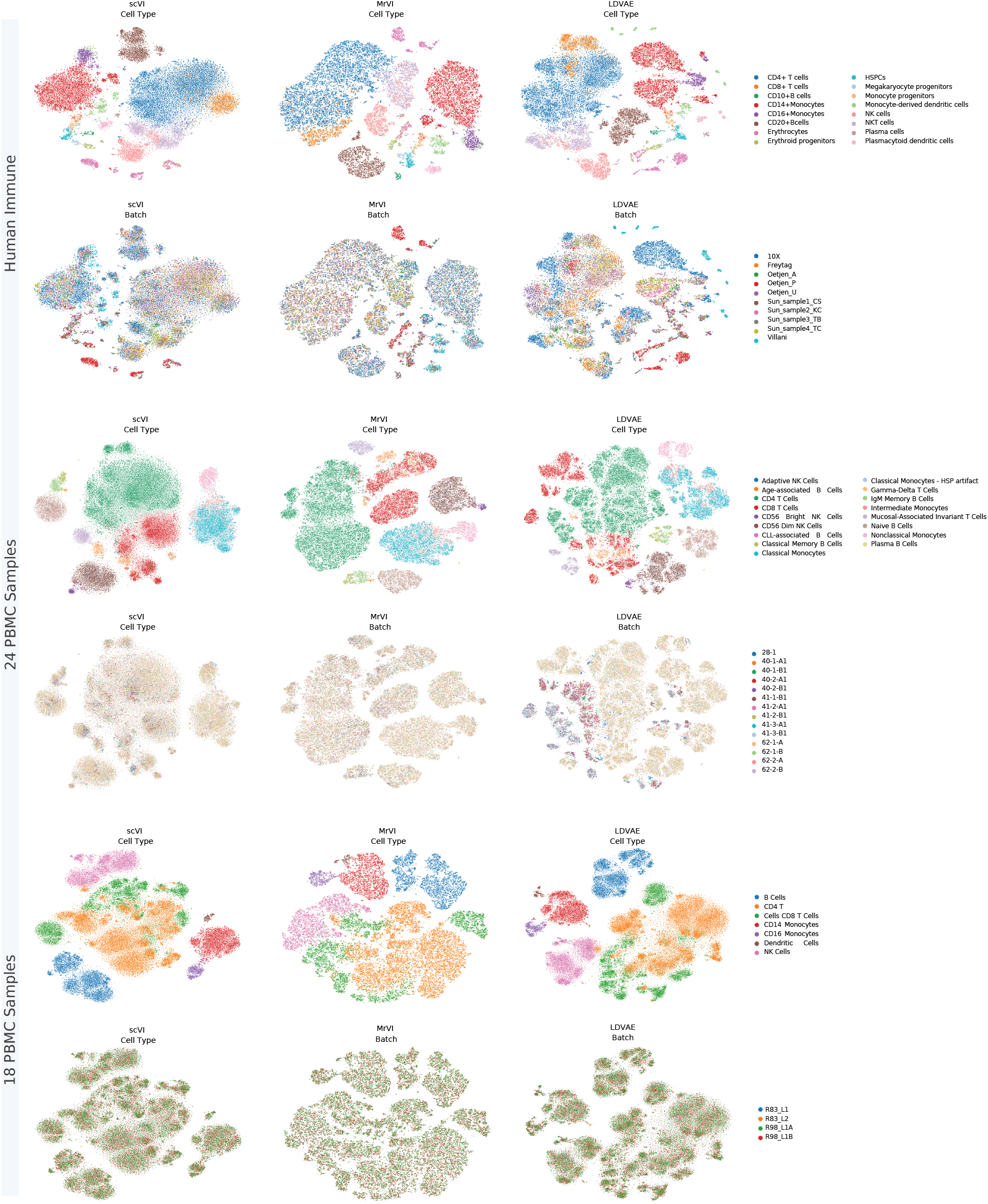
t-SNE visualization of integrated embeddings for each VAE-based model on immune datasets. Rows correspond to datasets (top to bottom): Human Immune (OpenProblems), 24 PBMC Samples (Zenodo 8020792), and 18 PBMC Samples (Zenodo 11100300). Columns show integration models (left to right): scVI, MrVI and LDVAE. For each dataset-model combination, the top panel shows the t-SNE embedding of the learned low-dimensional representation coloured by cell-type annotation, and the bottom panel shows the same embedding coloured by batch/sample identity (keys at right).

**Figure 7.**
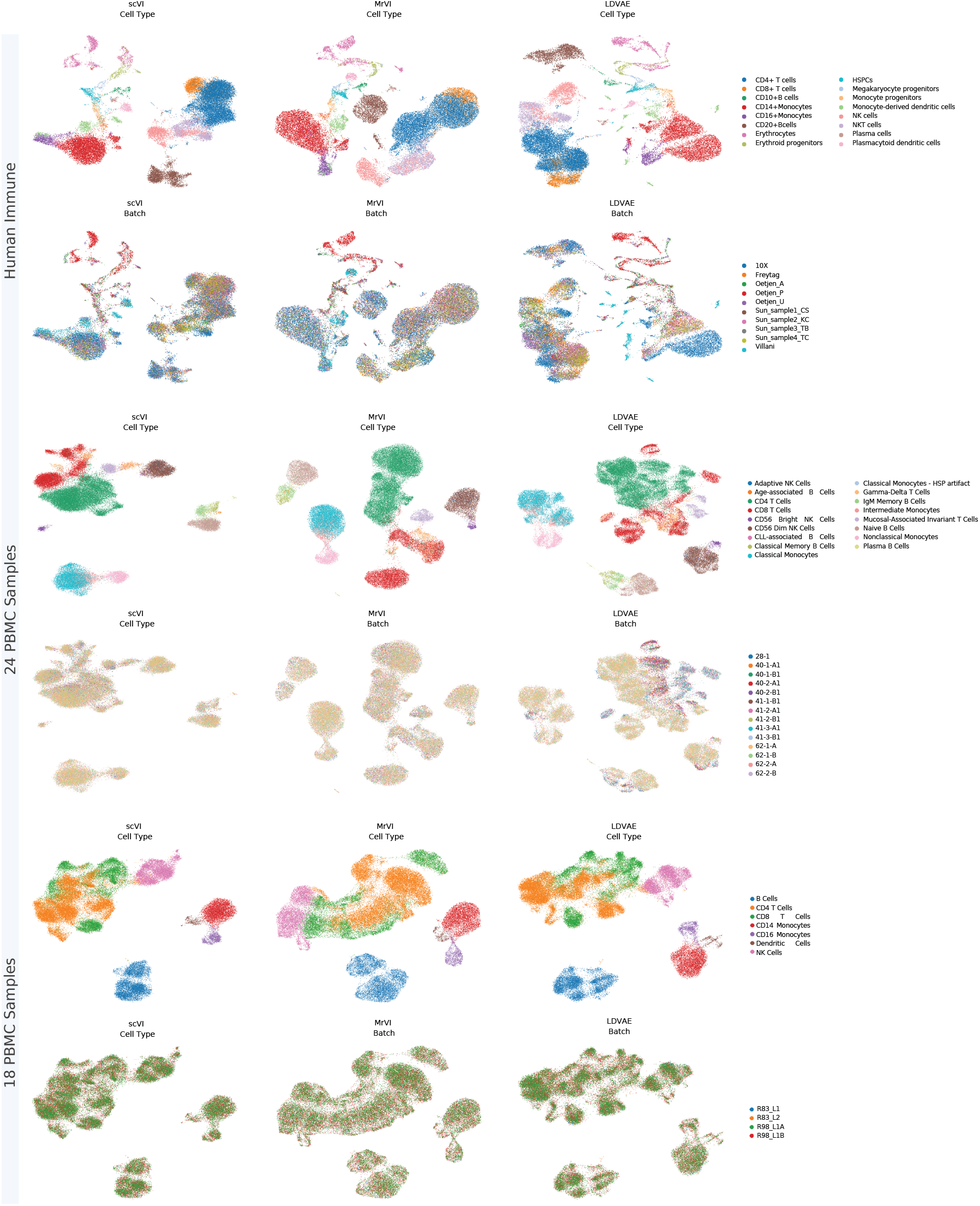
UMAP visualization of integrated embeddings for each VAE-based model on immune datasets. Rows correspond to datasets (top to bottom): Human Immune (OpenProblems), 24 PBMC Samples (Zenodo 8020792), and 18 PBMC Samples (Zenodo 11100300). Columns show integration models (left to right): scVI, MrVI and LDVAE. For each dataset-model combination, the top panel shows the UMAP embedding of the learned low-dimensional representation coloured by cell-type annotation, and the bottom panel shows the same embedding coloured by batch/sample identity (keys at right).

Figure 8 presents a t-SNE mosaic of the Zenodo 8020792 (24 PBMC samples) dataset using HVGs, illustrating how the embeddings change as all model hyperparameters are varied in a paired and crossed fashion. Similarly, Figure 9 shows the t-SNE mosaic for Human Immune and Figure 10 shows the t-SNE mosaic for Zenodo 11100300 (18 PBMC Samples). In Figure 8, embeddings show the separation of cell populations under different VAE model configurations (scVI, MrVI, LDVAE) across varying numbers of latent dimensions, hidden units, and layers. The first panel in this Figure shows scVI. We can observe that increasing latent dimensionality generally tightens the same cell-type clusters and increases the separation between phenotypes. The clearest improvements appear at higher latent sizes, with the best embeddings at *n*_latent_ = 50 for shallow networks (e.g., *n*_layers_ = 1 with *n*_hidden_ ∈{128, 256}). Moderate gains from increasing hidden width (128→ 256) are most visible in smaller latent sizes (around 20), while benefits are marginal in larger latent spaces. In contrast, deeper encoders (*n*_layers_ = 3) tend to fragment the embedding and reduce the gains of larger *n*_latent_, suggesting detrimental interactions between depth and latent capacity for this dataset.

**Figure 8.**
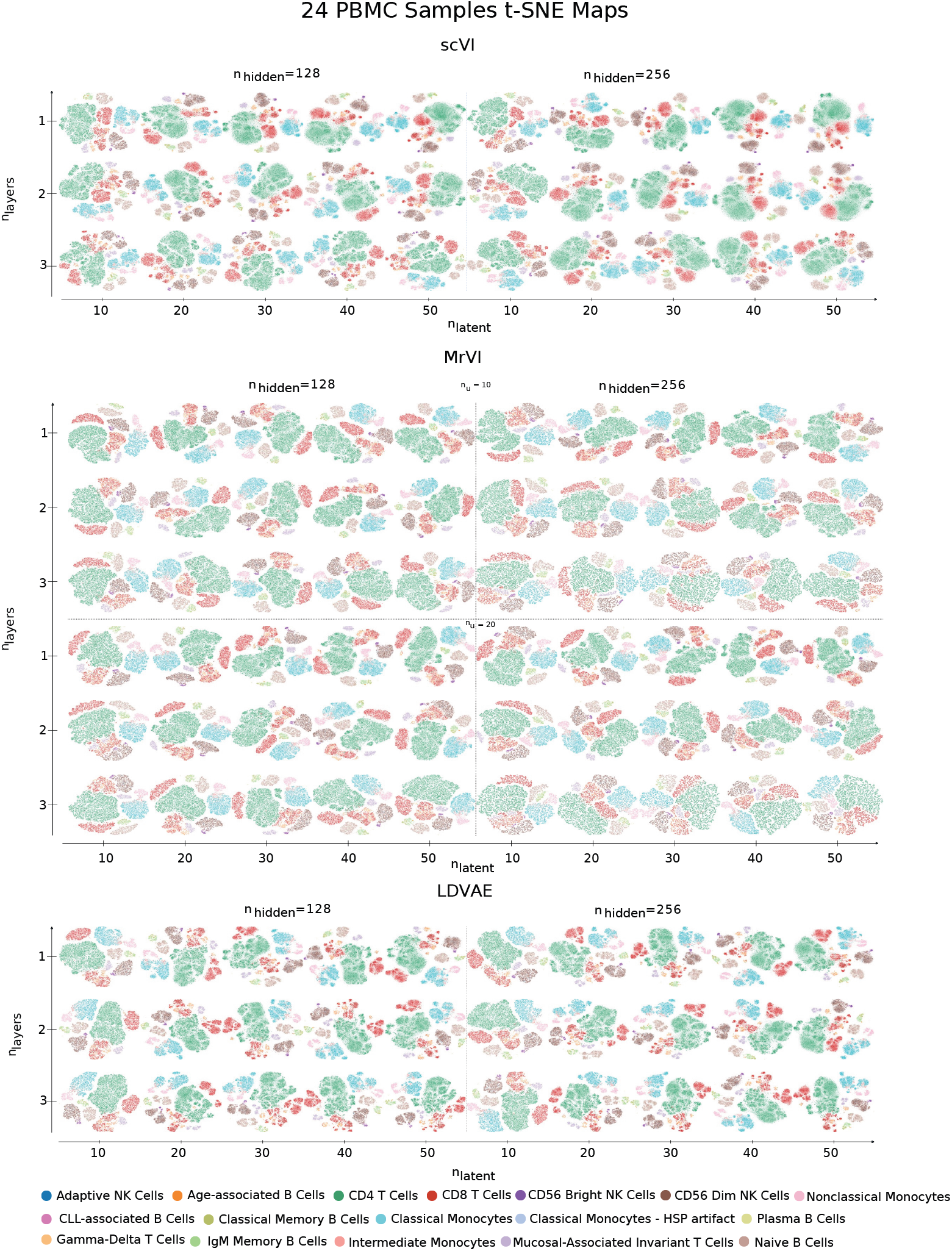
Architecture-dependent t-SNE maps for the 24 PBMC samples benchmark under HVG regime. t-SNE visualizations of the 24 PBMC Samples (Zenodo 8020792) computed from the learned latent embeddings of scVI (top), MrVI (middle) and LDVAE (bottom) after training on the top 5,000 HVG. Within each model family, mini-panels are arranged by shared latent dimensionality (nlatent = 10, 20, 30, 40, 50; horizontal), network depth (nlayers = 1–3; vertical) and hidden-layer width (nhidden = 128, left; 256, right). For MrVI, results are shown separately for the sample-aware latent dimension (nu = 10, upper block; nu = 20, lower block). Each point represents a cell and is coloured by cell-type annotation (keys at the bottom). This figure provides a qualitative assessment of how embedding geometry and cell-population separation vary across architectural settings.

**Figure 9.**
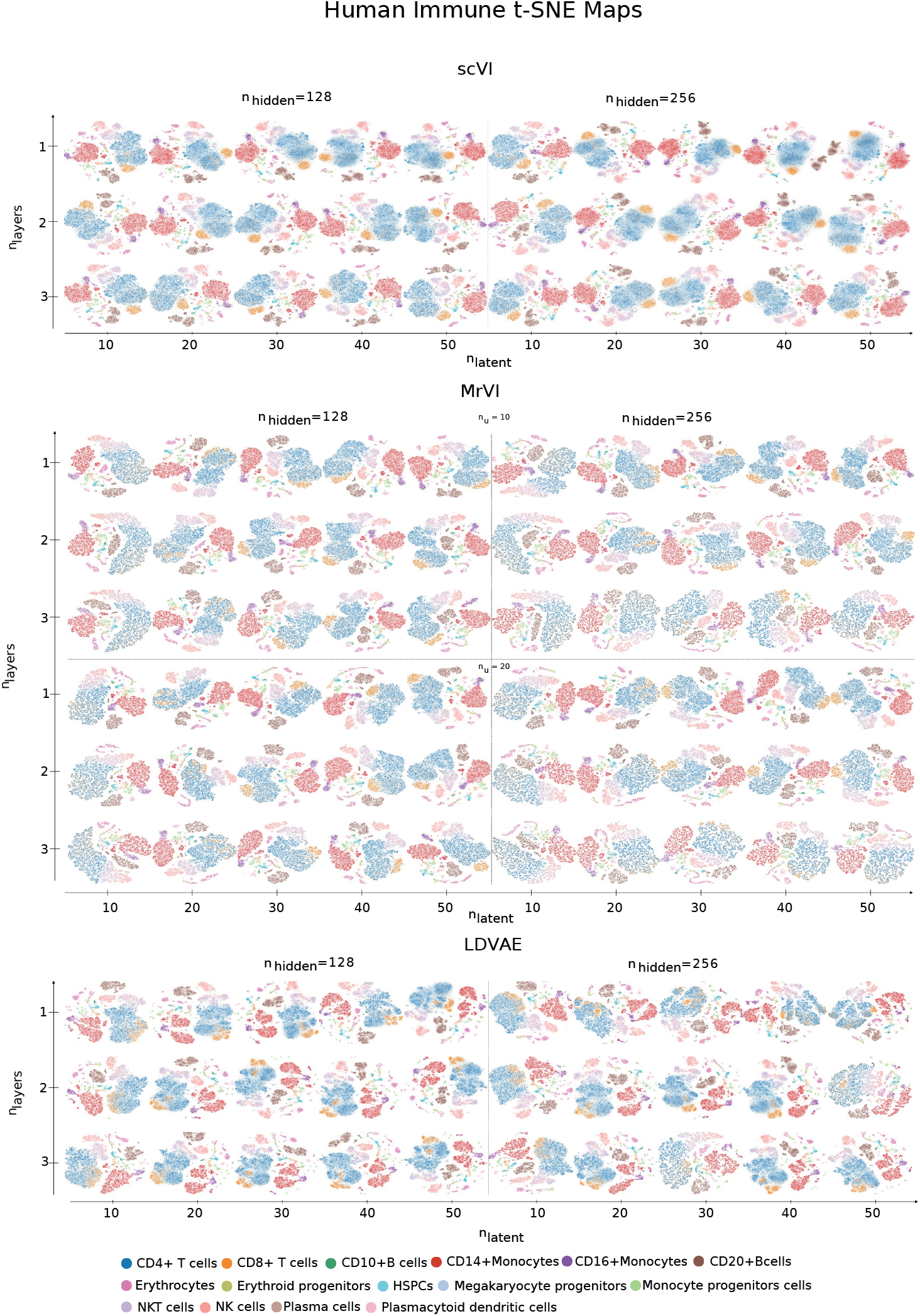
Architecture-dependent t-SNE maps for the Human Immune dataset benchmark under HVG regime. t-SNE visualizations of the Human Immune dataset (Open-Problems) computed from the learned latent embeddings of scVI (top), MrVI (middle) and LDVAE (bottom) after training on the top 5,000 HVG. Within each model family, mini-panels are arranged by shared latent dimensionality (nlatent = 10, 20, 30, 40, 50; horizontal), network depth (nlayers = 1–3; vertical) and hidden-layer width (nhidden = 128, left; 256, right). For MrVI, results are shown separately for the sample-aware latent dimension (nu = 10, upper block; nu = 20, lower block). Each point represents a cell and is coloured by cell-type annotation (keys at the bottom). This figure provides a qualitative assessment of how embedding geometry and cell-population separation vary across architectural settings.

**Figure 10.**
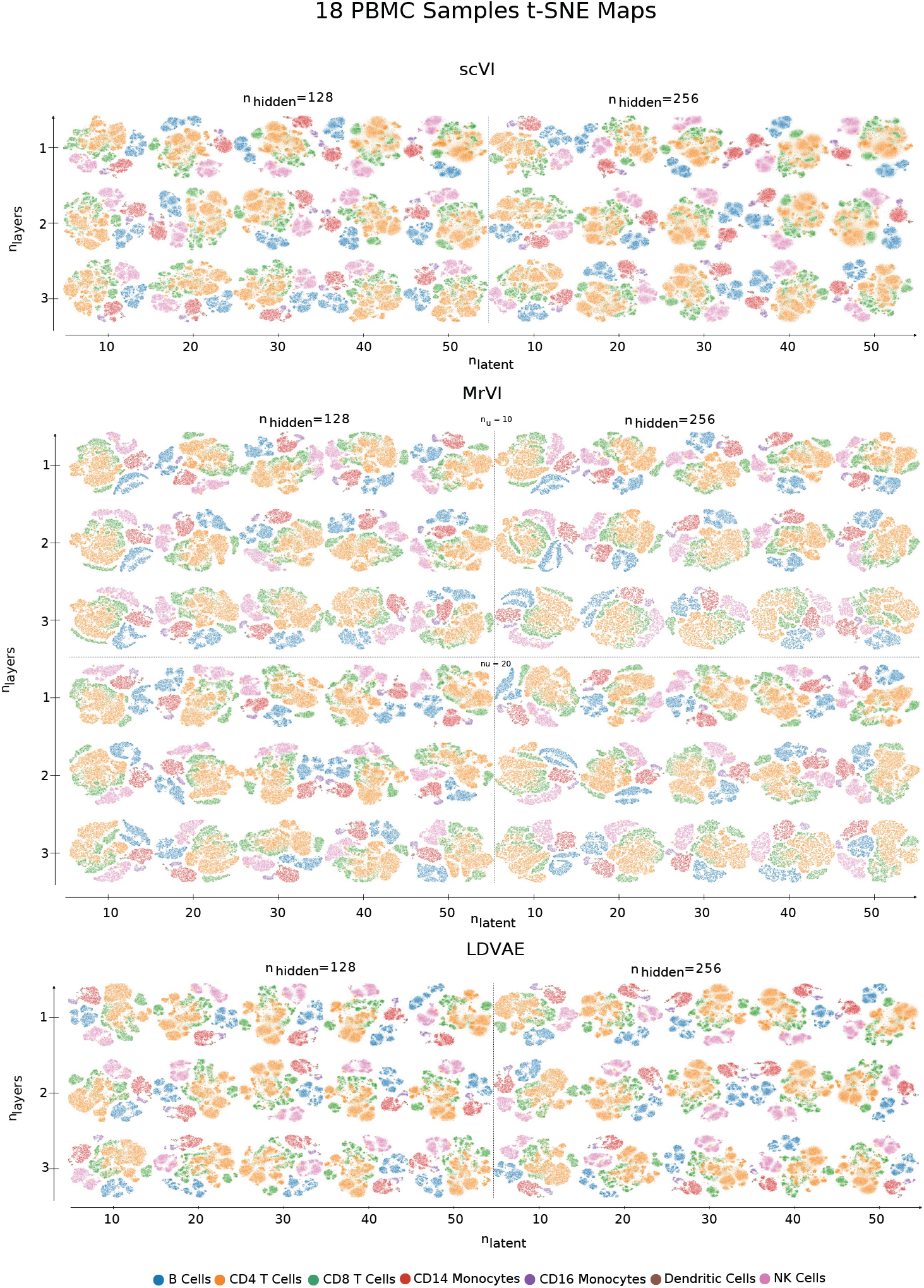
Architecture-dependent t-SNE maps for the 18 PBMC samples benchmark under HVG regime. t-SNE visualizations of the 18 PBMC Samples (Zenodo 11100300) computed from the learned latent embeddings of scVI (top), MrVI (middle) and LDVAE (bottom) after training on the top 5,000 HVG. Within each model family, mini-panels are arranged by shared latent dimensionality (nlatent = 10, 20, 30, 40, 50; horizontal), network depth (nlayers = 1–3; vertical) and hidden-layer width (nhidden = 128, left; 256, right). For MrVI, results are shown separately for the sample-aware latent dimension (nu = 10, upper block; nu = 20, lower block). Each point represents a cell and is coloured by cell-type annotation (keys at the bottom). This figure provides a qualitative assessment of how embedding geometry and cell-population separation vary across architectural settings.

In the second panel of Figure 8 we see the embeddings of MrVI. For MrVI, compact clustering typically emerges in the mid–latent regime (*n*_latent_ ≈20–40). Increasing from *u* = 10 to *u* = 20 yields improvements in select settings (e.g., *n*_hidden_ ∈{128, 256}, *n*_layers_ = 3, *n*_latent_ ∈{40, 50}), but this effect is not uniform across the grid. At *u* = 10, pushing *n*_latent_ > 40 can overspread clusters (e.g., *n*_hidden_ = 128, *n*_layers_ = 2, *n*_latent_ = 40 →50)), whereas *u* = 20 better preserves compactness at comparable latent sizes. Depth gives a little advantage and can modestly degrade separation, indicating MrVI is more sensitive to balancing *n*_latentU_ with *n*_latentZ_ than to adding layers.

In the third panel of Figure 8, we see the embeddings of LDVAE. In the case of LDVAE, it benefits most from increased hidden width. Raising *n*_latent_ from 10 to 50 improves separation, while adding the depth of the encoder (*n*_layers_ = 2 or 3) rarely changes the qualitative picture.

#### Per-Metric Performance on Human Immune - scVI

Table 6 reports a full hyperparameter sweep on the human immune dataset using scVI. There is a clear trade-off across *n*_hidden_ ∈ {128, 256}, *n*_latent_ ∈ {10, 20, 30, 40, 50}, *n*_layers_ ∈ {1, 2, 3}, and Features ∈ {Full, HVG}. Using the Overall score, the best configuration is 0.78105 at (256, 40, 2, HVG), while the worst is 0.73765 at (256, 10, 1, FULL). The batch-correction overall peaks under HVG with larger latent dimensionality (256, 50, 1, HVG), whereas the biological-conservation overall prefers HVG with smaller latent (256, 10, 1, HVG). These extremes indicate that model capacity and feature selection jointly set a spectrum between batch mixing and biological separability.

As shown in the cross-metric bar chart provided in Figure 16, changing *n*_hidden_ from 128 to 256 showed in general small gains in batch-oriented metrics, Batch ASW (30/30), iLISI (22/30), Overall (batch) (23/30), and connectivity (28/30) improved in most paired settings, while PCR is mixed (16/30 up). Biological/label metrics often decrease (e.g., Overall(bio) 11/30 up, NMI 11/30 up, Label ASW 2/30 up). Increasing *n*_layers_ from 1 to 3 further tilts toward batch mixing, with PCR (19/20) and iLISI (16/20) improving in most paired settings and overall batch success in 11/20 cases. The trajectory also improves in 19/20 pairs. In contrast, clustering agreement declines (NMI 3/20 up, 17/20 down; ARI 9/20 up, 11/20 down), and the biological overall decreases in 14/20 pairs. Other label metrics are mixed: The ASW label and the isolated ASW label more often increase (14/20 and 13/20, respectively), but the isolated label F1 worsens in 20/20 cases. The Overall score shifts only slightly (11/20 up), consistent with a small net effect at the aggregate level. HVG outperforms FULL on average for overall, batch-correction metrics, and clustering agreement (notably ARI), with only small penalties in connectivity and trajectory and near-neutral effects on label ASW; HVG is therefore a strong default.

To quantify the latent effect, we performed paired comparisons while holding (*n*_hidden_, *n*_layers_, Features) fixed and stepping *n*_latent_ through 10→ 20→ 30 →40 →50 (4 steps ×12 fixed triplets = 48 stepwise comparisons per metric). Batch-correction metrics improve in the large majority of steps: Overall(batch) 37/48 up, Batch ASW 37/48 up, PCR 33/48 up, and iLISI 35/48 up. Clustering agreement also trends upward (NMI 27/48 up; ARI 28/48 up). Label-compactness metrics tend to soften (Label ASW 14/48 up, 34/48 down; Isolated Label ASW 16/48 up, 32/48 down), and trajectory shows a near balance (23/48 up, 25/48 down). The biological overall is mixed on a stepwise basis (24/48 up, 24/48 down), yet the endpoint comparison (*n*_latent_=50 versus 10) improves in 10/12 fixed triplets. The Overall rises in 31/48 steps, and the endpoint 10→50 improves in 11/12 cases.

#### Per-Metric Performance on Human Immune - MrVI

Table 3 reports the MrVI sweep on the human immune dataset across *n*_hidden_ ∈ {128, 256}, *n*_latent_ ∈{10, 20, 30, 40, 50}, *n*_layers_ ∈ {1, 2, 3}, *n*_latent_u_ ∈ {10, 20},and Features ∈ {Full, HVG}. Using the Overall score,the best configuration is 0.76041 at (256, 40, 1, *u*=20, Full), while the worst is 0.66557 at (128, 10, 3, *u*=20, HVG). The same (256, 40, 1, *u*=20, Full) setting also sits at or near the top for the batch and biology Overalls alongside NMI and ARI, indicating a pronounced peak. In contrast, label fidelity and temporal continuity reach their best values at more conservative capacity: Label ASW near (256, 30, 2, *u*=20, HVG) and Trajectory near (128, 10, 2, *u*=10, Full).

**Table 3.**
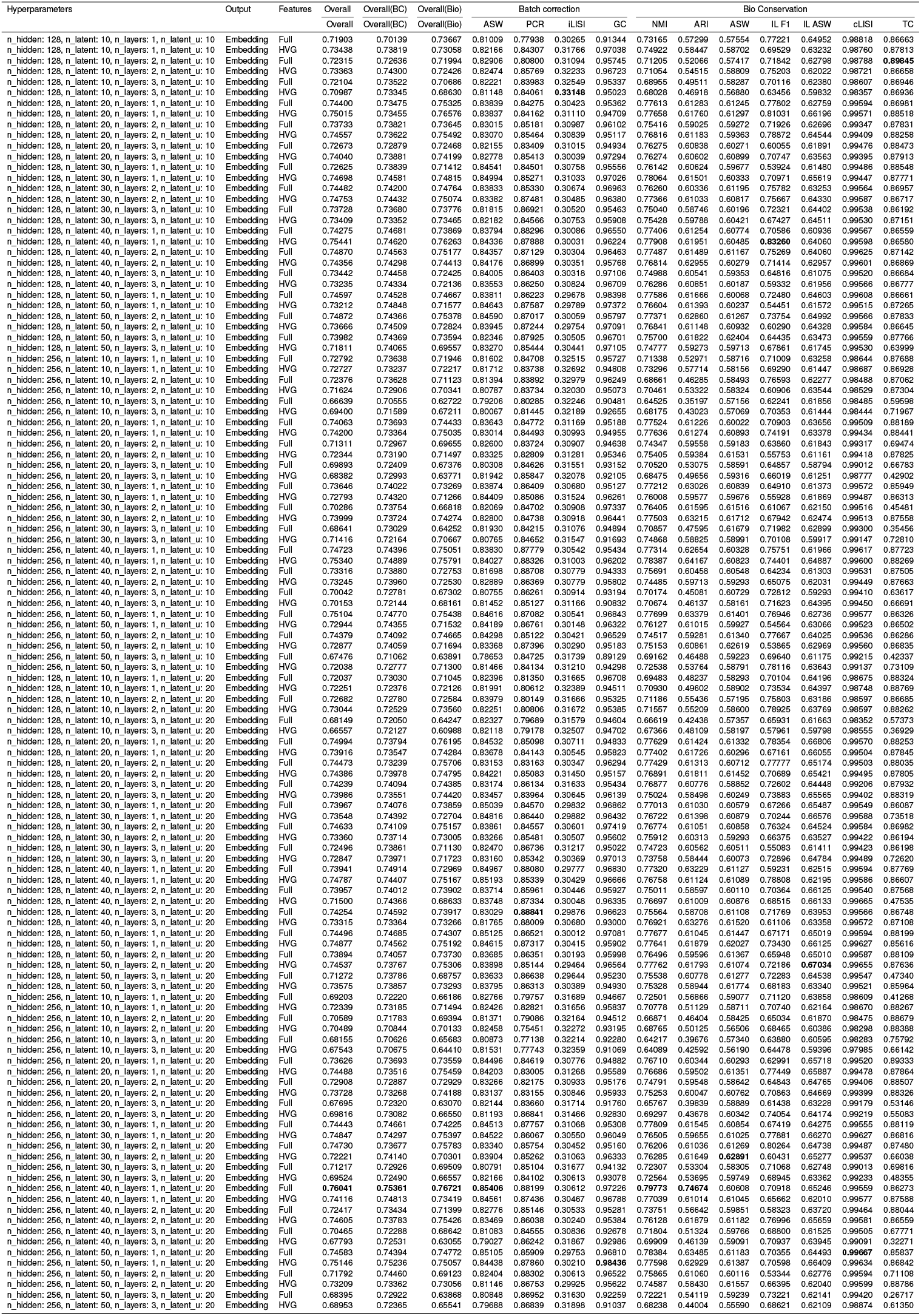
Performance of MrVI on the Human Immune Dataset.

As shown in the cross-metric bar chart provided in Figure 17, changing hidden size from 128 to 256 generally depresses the aggregate metrics in this MRVI grid. Across matched pairs that fix (*n*_latent_, *n*_layers_, *u*, Features), Overall rises in 17*/*60 and falls in 43*/*60; the batch Overall rises in 13*/*60 and falls in 47*/*60; the biology Overall rises in 13*/*60 and falls in 47*/*60. Batch ASW drops in 48*/*60, NMI in 48*/*60, ARI in 44*/*60, and Graph Connectivity in 44*/*60 pairs. The main counter-trend is iLISI, which improves in 48*/*60 when moving to 256. Trajectory is mixed (26*/*60 up, 34*/*60 down). Overall, *n*_hidden_=128 is the safer default unless iLISI is prioritized.

Increasing depth from *n*_layers_=1 to 3 also pushes downward on the composite metrics. Over matched pairs holding (*n*_hidden_, *n*_latent_, *u*, Features) fixed, Overall rises in 4*/*40 and falls in 36*/*40; the batch Overall rises in 4*/*40 and falls in 36*/*40; the biology Overall rises in 3*/*40 and falls in 37*/*40. Batch ASW decreases in 37*/*40 and Graph Connectivity in 32*/*40; clustering agreement weakens (NMI up 1*/*40, down 39*/*40; ARI up 12*/*40, down 28*/*40). iLISI improves in 32*/*40 and PCR in 12*/*40. In short, deeper models tend to inflate iLISI but hurt agreement, connectivity, and the composite Overalls; shallow depth (1) is preferred for balance.

Latent dimensionality in *z* is the strongest and most favorable driver. Endpoint comparisons (50 vs. 10) at fixed (*n*_hidden_, *n*_layers_, *u*, Features) show Overall improving in 23*/*24 cases, the batch Overall in 24*/*24, and the biology Overall in 20*/*24. NMI and ARI each improve in 24*/*24, Graph Connectivity in 21*/*24, Label ASW in 23*/*24, and Isolated-Label ASW in 19*/*24. The main downside is Trajectory, which falls in 18*/*24 endpoints. Stepwise behavior is not strictly monotone for every metric, but the end state at high latent (40–50) is decisively better on aggregate and agreement metrics. Practically, pushing *n*_latent_ high with shallow depth is the winning recipe for the composite scores without sacrificing label compactness in this MrVI setting.

The unshared *u* latent shows nuanced tradeoffs. Moving from *u*=10 to *u*=20 at fixed (*n*_hidden_, *n*_latent_, *n*_layers_, Features), Overall is up in 27*/*60 and down in 33*/*60; the batch Overall is up in 25*/*60 and down in 34*/*60; the biology Overall is up in 27*/*60 and down in 33*/*60. Several structure and label metrics tilt positive: Batch ASW improves in 40*/*60, Label ASW in 36*/*60, Isolated-Label ASW in 43*/*60, and Trajectory in 33*/*60 pairs; PCR tends to fall (24*/*60 up, 36*/*60 down), and Connectivity is roughly balanced (27*/*60 up, 33*/*60 down). Thus, *u*=20 aids label compactness and Batch ASW, while composite Overalls move only modestly.

Feature selection has a small but broad positive tilt for HVG. Pairing HVG against Full at fixed (*n*_hidden_, *n*_latent_, *n*_layers_, *u*), Overall is higher in 32*/*60 pairs (lower in 28*/*60), ARI in 36*/*60, NMI in 31*/*60, iLISI in 33*/*60, and Trajectory tilts higher on average. Effects are modest in magnitude but consistent enough that HVG is a reasonable default; the single global optimum here, however, happens to be Full at (256, 40, 1, *u*=20).

#### Per-Metric Performance on Human Immune - LDVAE

Table 4 summarizes the sweep over *n*_hidden_ ∈{128, 256}, *n*_latent_ ∈{10, 20, 30, 40, 50}, *n*_layers_ ∈{1, 2, 3,} and Features ∈{Full, HVG} for LDVAE. Using the Overall score, the best configuration is 0.75774 at (*n*_hidden_=128, *n*_latent_=30, *n*_layers_=2, HVG), while the worst is 0.61916 at (256, 50, 2, Full). The batch-correction overall peaks at (256, 40, 2, HVG) and the biological overall coincides with the Overall winner at (128, 30, 2, HVG). Individual metric optima are spread: Batch ASW is highest at (128, 50, 1, HVG); PCR at (128, 30, 2, HVG); iLISI at (256, 10, 3, HVG); graph connectivity at (128, 10, 1, HVG); NMI and ARI at (128, 40, 2, HVG); Label ASW at (256, 10, 3, HVG); trajectory at (256, 30, 1, Full).

**Table 4.**
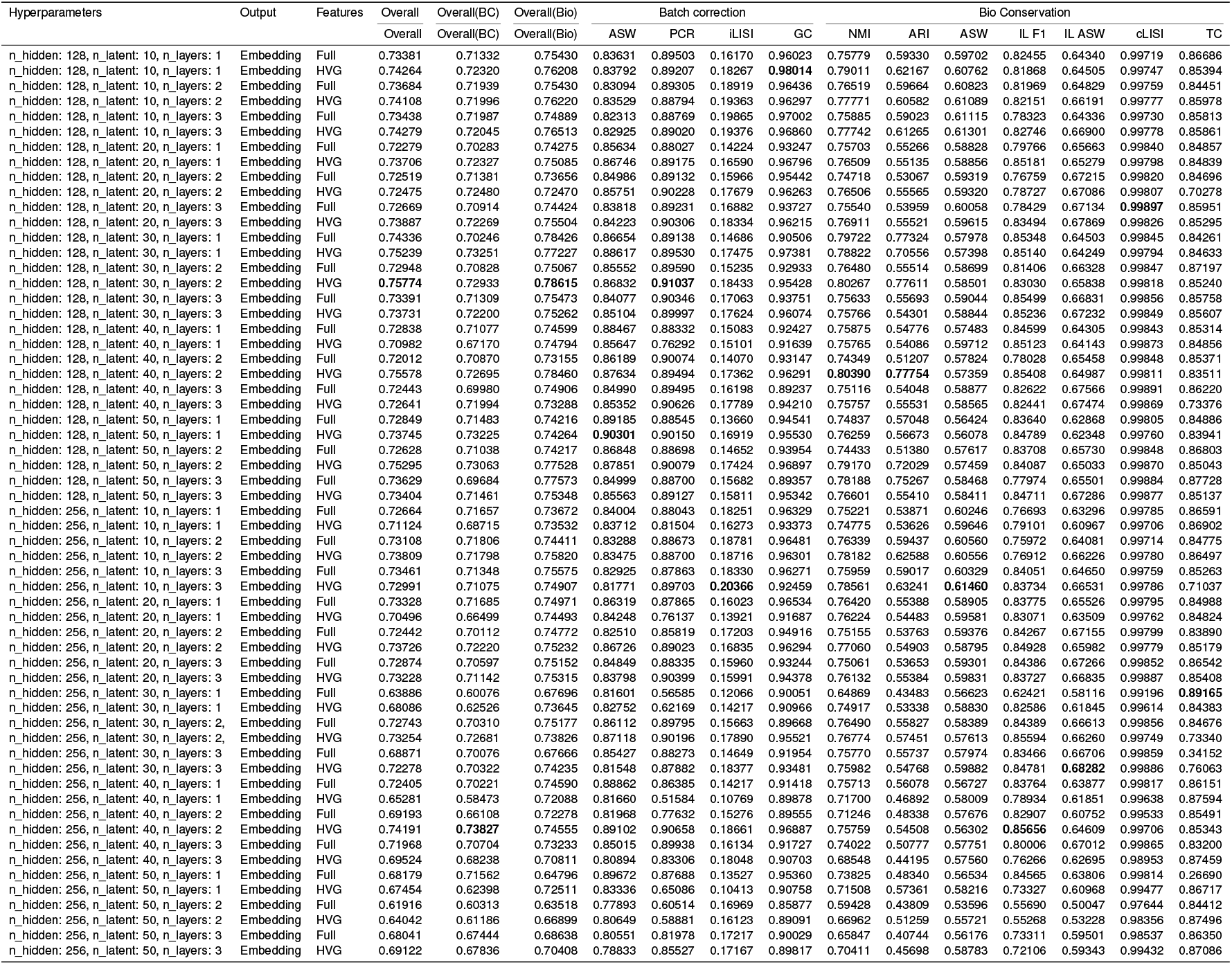
Performance of LDVAE on the Human Immune Dataset.

As shown in the cross-metric bar chart provided in Figure 18, changing hidden size from 128 to 256 generally reduces the composites and many structure metrics in this grid. Across paired settings that fix (*n*_latent_, *n*_layers_, Features), Overall improves in 4*/*30 and worsens in 26*/*30 pairs; the batch overall in 5*/*30 up and 25*/*30 down; the biology overall in 6*/*30 up and 24*/*30 down. By metric, Batch ASW is up in 13*/*30, PCR in 5*/*30, iLISI in 12*/*30, connectivity in 10*/*30, NMI in 9*/*30, ARI in 9*/*30, Label ASW in 11*/*30, and trajectory in 16*/*30 pairs. The net effect is that moving to 256 tends to hurt the composites and clustering agreement, with trajectory the only metric showing a near-balanced tilt.

Increasing depth from *n*_layers_=1 to 3 is net favorable here for the composite scores, while shifting the detailed balance of batch versus biology. Over paired settings fixing (*n*_hidden_, *n*_latent_, Features), Overall rises in 13*/*20 and falls in 7*/*20; the batch overall 11*/*20 up, 9*/*20 down; the biology overall 12*/*20 up, 8*/*20 down. PCR and iLISI improve strongly (15*/*20 and 18*/*20 up, respectively), Label ASW improves in 17*/*20, and trajectory in 12*/*20. In contrast, Batch ASW drops in 19*/*20, while NMI and ARI are more often down (8*/*12 down for both). Thus depth promotes certain batch harmonization statistics (PCR, iLISI) and label compactness, with mild trajectory gains, but it softens Batch ASW and clustering agreement.

Latent dimensionality in *z* does not help in this dataset when pushed to the high end. Endpoint comparisons between *n*_latent_=50 and 10 at fixed (*n*_hidden_, *n*_layers_, Features) show Overall up in only 2*/*12 and down in 10*/*12; the batch overall 3*/*12 up, 9*/*12 down; the biology overall 2*/*12 up, 10*/*12 down. Endpoints for individual metrics show Batch ASW 7*/*12 up, but PCR 3*/*12 up, iLISI 0*/*12 up, connectivity 1*/*12 up, NMI 2*/*12 up, ARI 3*/*12 up, Label ASW 0*/*12 up, trajectory 5*/*12 up. Stepwise trends (10→20→30→40→50) corroborate this: Batch ASW rises in 35*/*48 steps, but many other metrics tilt downward by the time we reach 50. In short, large *n*_latent_ is not advantageous in this sweep; moderate latent (around 30) with shallow-to-moderate depth is where the composite and biology overalls peak.

Comparing HVG to Full at matched (*n*_hidden_, *n*_latent_, *n*_layers_) pairs, HVG gives consistent, modest gains in the composites and agreement metrics: up in 21*/*30 pairs, batch overall (22*/*30), biology overall (19*/*30). For individual metrics, HVG improves iLISI (21*/*30 up), connectivity (19*/*30), NMI (22*/*30), ARI (19*/*30), and Label ASW (15*/*30 up with many near-ties). PCR averages slightly down despite more ups than downs, indicating a few large negative shifts. Overall, HVG remains a sensible default here given its small but broad gains.

#### Per-Metric Performance on ZENODO 8020792 (24 PBMC SAMPLES) - scVI

Table 5 reports the scVI sweep on Zenodo 8020792 over *n*_hidden_ ∈{128, 256}, *n*_latent_ ∈{10, 20, 30, 40, 50}, *n*_layers_ ∈{1, 2, 3}, and Features∈ {Full, HVG}. The best Overall is 0.67480 at (256, 50, 1, HVG), while the worst is 0.58505 at (128, 10, 2, Full). Batch-correction overall peaks at (256, 50, 2, HVG), whereas the biology overall reaches its maximum at (256, 20, 1, Full). By individual metrics, the strongest batch mixing occurs at (256, 50, 1, HVG) for Batch ASW (0.93546), PCR (0.80196), and iLISI (0.31896), graph connectivity peaks at(256, 50, 1, Full) (0.88811), clustering agreement peaks at (256, 20, 1, Full) for NMI (0.84489) and at (128, 50, 2, HVG) for ARI (0.86089), while label compactness (Label ASW) is highest at (128, 10, 1, HVG) (0.54473). The trajectory score is not applicable here (N/A) because the dataset lacks the necessary trajectory information.

**Table 5.**
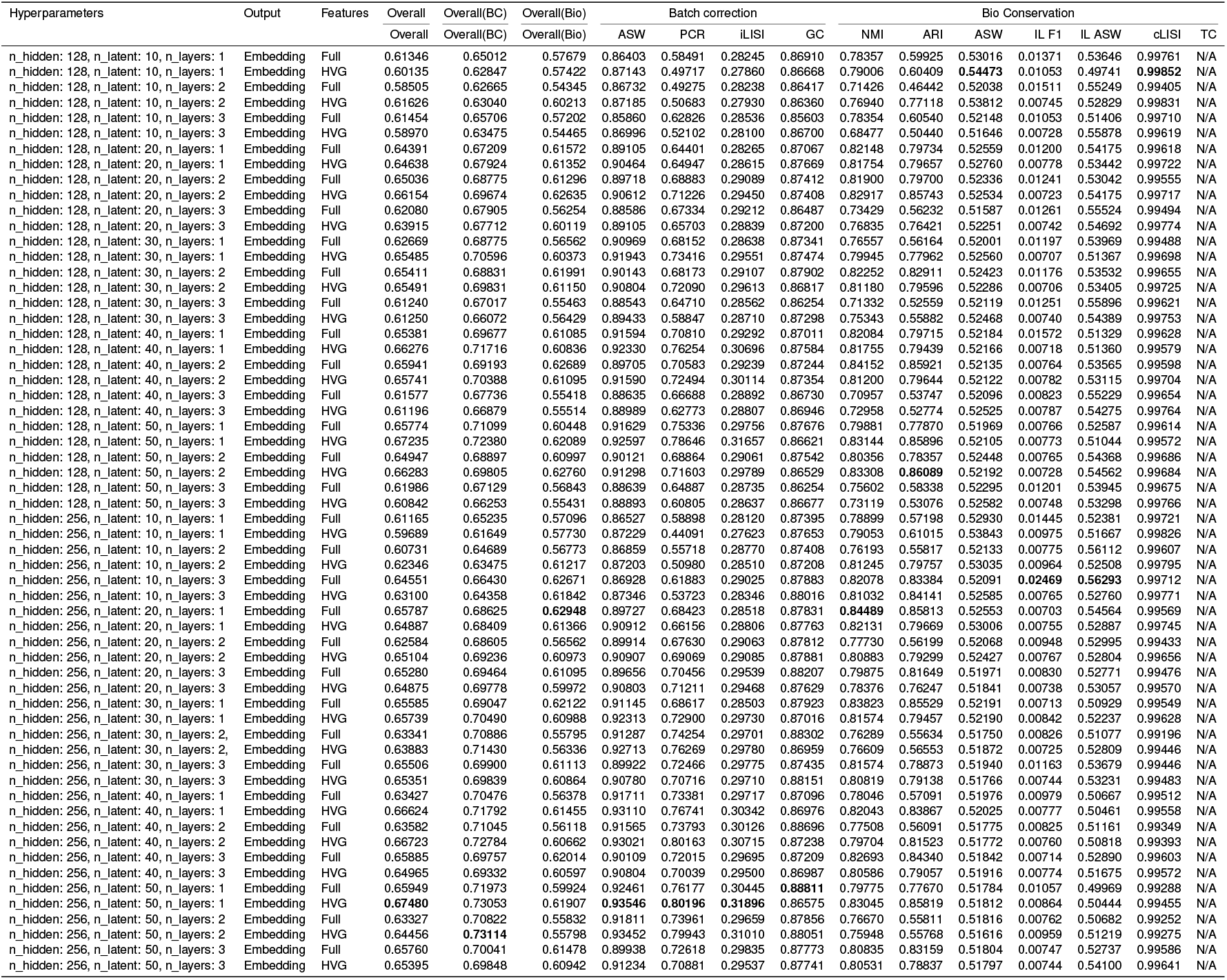
Performance of scVI on the Zenodo 8020792 (24 PBMC Samples) Dataset.

As shown in the cross-metric bar chart provided in Figure 19, increasing hidden size from 128 to 256 improves overall performance in 20*/*30 matched pairs, the batch overall in 26*/*30, and the biology overall in 17*/*30. Increasing depth from *n*_layers_=1 to 3 is generally unfavorable for the composites: Overall rises in 4*/*20 and falls in 16*/*20 pairs; the batch overall is 8*/*20 up and 12*/*20 down; the biology overall is 4*/*20 up and 16*/*20 down. In contrast, increasing latent dimensionality has the most systematic positive effect on batch correction: stepwise 10→ 20→ 30→ 40→ 50 comparisons show an overall increase in 34*/*48 steps and batch overall up in 39*/*48; Batch ASW and iLISI rise in 41*/*48 and 38*/*48 steps, respectively. The biology overall is mixed stepwise (23*/*48 up, 25*/*48 down), and the endpoint comparison 50− 10 is strongly positive for Overall (12*/*12 up) and batch overall (12*/*12up) while only modestly positive for the biology overall (7*/*12 up). Finally, HVG outperforms Full: HVG− Full results in Overall (18*/*30 up), batch overall (17*/*30 up), and biology overall (15*/*30 up), with additional gains often seen in NMI/ARI and connectivity.

#### Per-Metric Performance on ZENODO 8020792 (24 PBMC SAMPLES) - MrVI

Table 6 reports the MrVI sweep on Zenodo 8020792 over *n*_hidden_ ∈{128, 256}, *n*_latent_ ∈{10, 20, 30, 40, 50},*n*_layers_ ∈{1, 2, 3}, *n*_latent_u_ ∈{10, 20} and Features ∈{Full, HVG}. The best Overall is 0.62984 at (256, 40, 1, Full), while the worst is 0.54528 at (256, 10, 3, HVG).

**Table 6.**
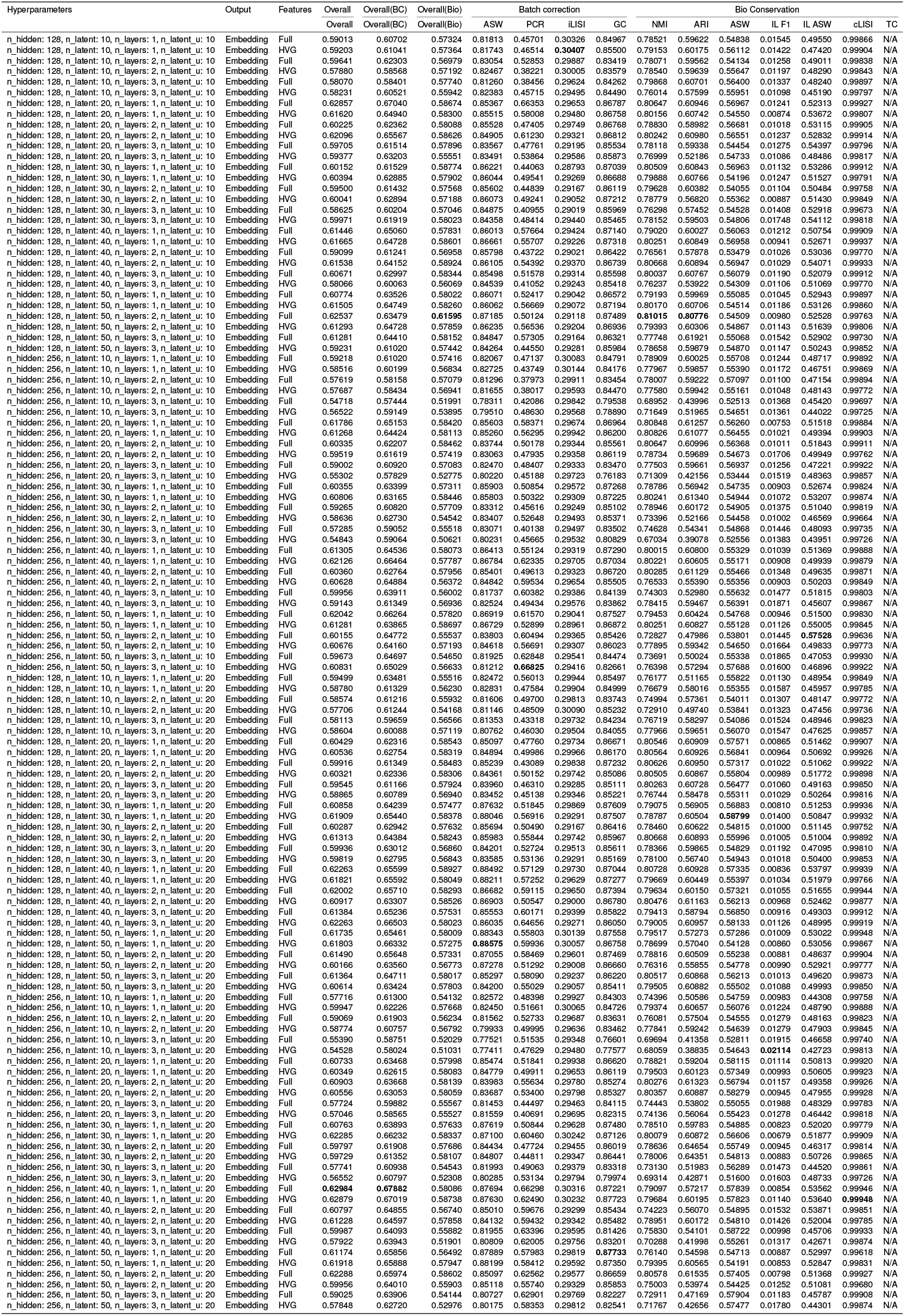
Performance of MrVI on the Zenodo 8020792 (24 PBMC Samples)

As shown in the cross-metric bar chart provided in Figure 20, changing hidden size from 128 to 256 tends to lower the composite scores in this grid. Across matched pairs that fix (*n*_latent_, *n*_layers_, Features), Overall improves in 9*/*30 and worsens in 21*/*30 pairs; the batch overall in 9*/*30 up and 21*/*30 down; the biology overall in 8*/*30 up and 22*/*30 down. Metric-wise, moving to 256 often improves PCR and iLISI (many ups) and frequently raises isolated-label F1, but it more often reduces Batch ASW, graph connectivity, and clustering agreement (NMI/ARI), with label-compactness metrics showing mixed behavior. In short, larger hidden size favors certain batch harmonization statistics but usually hurts the aggregate objectives on this dataset.

Increasing depth from *n*_layers_=1 to 3 is broadly unfavorable for the composites. In matched pairs, Overall rises in 1*/*20 and falls in 19*/*20; the batch overall 1*/*20 up and 19*/*20 down; the biology overall 3*/*20 up and 17*/*20 down. Batch ASW and graph connectivity decrease in nearly all pairs, iLISI and PCR improve only occasionally, and clustering agreement (NMI/ARI) more often declines. The consistent exception is isolated-label F1, which increases in all pairs. Practically, depth erodes batch mixing and graph cohesion here and does not pay off in the aggregate.

The latent dimension shows interactions with capacity and features, but in this table the exact stepwise trends (10→50) cannot be paired systematically across fixed (*n*_hidden_, *n*_layers_, Features), so we avoid over-generalization. The best Overall occurs at moderate-to-high latent (40) with (256, 1, Full), while several batch and structure metrics peak at higher latent values in specific settings, illustrating that latent interacts with both hidden size and feature selection.

Sweeping *n*_latent_u_ causes the primary Overall increases in 37*/*60 pairs, the batch composite Overall(BC) in 40*/*60, while the biology composite Overall(Bio) decreases in aggregate (up 23*/*60, down 37*/*60). Batch metrics are the main beneficiaries: PCR rises in 39*/*60, iLISI in 40*/*60, and Batch ASW in 34*/*60. Graph connectivity slightly declines on average (up 23*/*60). For clustering agreement, NMI and ARI tilt down overall (NMI up 23*/*60; ARI up 24*/*60). Label compactness is mixed: Label ASW improves (up 37*/*60), whereas Isolated Label ASW and Isolated Label F1 tend to fall (up 20*/*60; up 26*/*60). cLISI is essentially flat (up 32*/*60).

Comparing feature selection at matched (*n*_hidden_, *n*_latent_, *n*_layers_), Full has a slight edge on the composite Overalls for this dataset: HVG minus Full shows small negative mean differences for Overall and the batch overall, whereas the biology overall is roughly balanced to slightly positive for HVG. However, we consider these differences negligible and still recommend HVG as a more favarobale option.

#### Per-Metric Performance on ZENODO 8020792 (24 PBMC SAMPLES) - LDVAE

Table 7 summarizes the LDVAE sweep on Zenodo 8020792 over *n*_hidden_ ∈{128, 256},*n*_latent_ ∈{10, 20, 30, 40, 50}, *n*_layers_ ∈{1, 2, 3}, and Features ∈{Full, HVG}. The best Overall is 0.60260 at (256, 50, 1, HVG), while the worst is 0.49913 at (256, 10, 2, Full). Across matched pairs, as shown in the cross-metric bar chart provided in Figure 21, increasing hidden size from 128 to 256 is favorable for LDVAE on this dataset. Overall improves in 19*/*30 pairs (down in 11*/*30), the batch-correction overall improves in 17*/*30 (down in 13*/*30), and the biology overall improves in 20*/*30 (down in 10*/*30). Thus, larger hidden units tend to raise all three composites here.

**Table 7.**
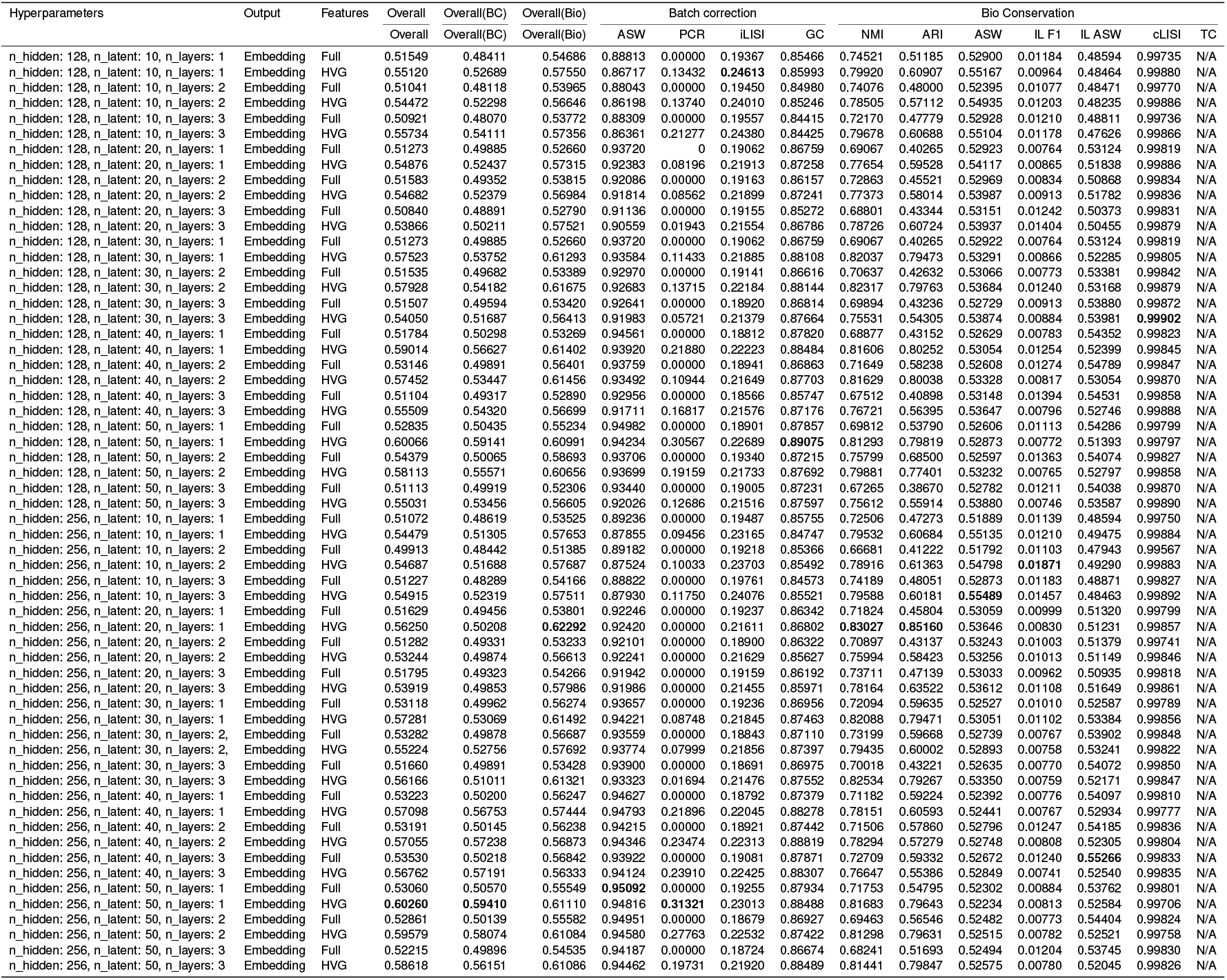
Performance of LDVAE on the Zenodo 8020792 (24 PBMC Samples)

Depth has the opposite effect. Comparing *n*_layers_=3 versus 1 at fixed (*n*_hidden_, *n*_latent_, Features), Overall rises in 6*/*20 and falls in 14*/*20 pairs; the batch-correction overall rises in 4*/*20 and falls in 16*/*20; the biology overall rises in 6*/*20 and falls in 14*/*20. Deeper encoders are broadly unfavorable for the aggregates in this benchmark.

Latent dimensionality shows the most systematic trend. Stepwise comparisons 10→20→30→40→50 aggregated over fixed (*n*_hidden_, *n*_layers_, Features) yield Overall up in 31*/*48 steps (down 16*/*48), the batch-correction overall up in 36*/*48 (down 11*/*48), and the biology overall up in 24*/*48 (down 23*/*48). The endpoint comparison50− 10 strengthens this picture: Overall is higher in 11*/*12 matched triplets (down 1*/*12), batch overall in 11*/*12 (down 1*/*12), and biology overall in 10*/*12 (down 2*/*12). Increasing the latent dimension therefore helps LDVAE’s aggregate scores here, especially the batch-oriented composite. Features strongly favor HVG for LDVAE in this dataset.

In matched (*n*_hidden_, *n*_latent_, *n*_layers_) pairs, HVG− Full for overall with counts 30*/*30 up, the batch overall 30*/*30 up, and the biology overall with 29*/*30 up. HVG consistently dominates Full for the composites.

Metric-wise optima align with these trends. The batch-correction metrics favor high latent and often larger hidden size: Batch ASW peaks at (256, 50, 1, Full), PCR at (256, 50, 1, HVG), while iLISI achieves its maximum at (128, 10, 1, HVG). Graph connectivity peaks at (128, 50, 1, HVG). Clustering agreement is strongest at moderate latent with higher hidden size, with NMI and ARI both maximized at (256, 20, 1, HVG). Label compactness (Label ASW) peaks at (256, 10, 3, HVG), reflecting that deeper and lower-latent settings can tighten local label structure even as they hurt the composite scores.

#### Per-Metric Performance on ZENODO 11100300 (18 PBMC SAMPLES) - scVI

Table 8 reports the scVI sweep on Zenodo 11100300 over *n*_hidden_ ∈{128, 256}, *n*_latent_∈{ 10, 20, 30, 40, 50}, *n*_layers_ ∈{1, 2, 3}, and Features ∈{Full, HVG}. The best Overall is 0.78615 at (256, 30, 1, HVG), while the worst Overall is 0.69409 at (256, 40, 1, HVG). The trajectory, isolated label F1, and isolated label ASW are not applicable for this dataset and therefore reported as N/A.

**Table 8.**
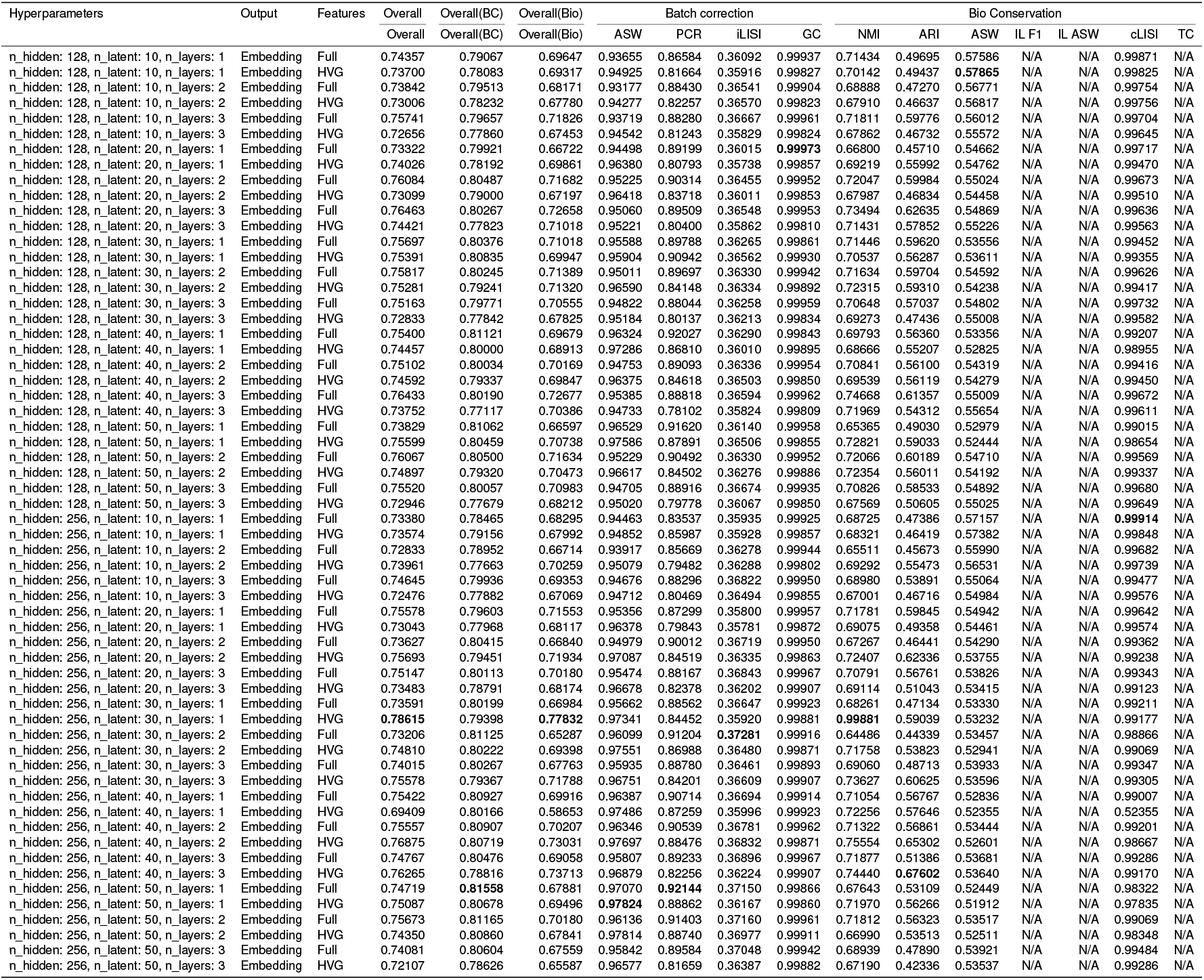
Performance of scVI on the Zenodo 11100300 (18 PBMC Samples)

As shown in the cross-metric bar chart provided in Figure 22, changing the hidden size from 128 to 256 consistently benefits batch correction but tends to reduce biology and the combined overall. In matched pairs that fix (*n*_latent_, *n*_layers_, Features), Overall improves in 10*/*30 and declines in 20*/*30 pairs, Overall(BC) improves in 20*/*30 and declines in 10*/*30, while Overall(Bio) improves in 10*/*30 and declines in 20*/*30. This indicates that a larger hidden size strengthens batch mixing on this dataset, but generally softens biology and the combined overall.

Depth primarily trades batch for biology with little net effect on the combined overall. Comparing *n*_layers_=3 against 1 at fixed (*n*_hidden_, *n*_latent_, Features), Overall rises in 9*/*20 and falls in 11*/*20 pairs (effectively flat). The batch-oriented Overall(BC) rises in 6*/*20 and falls in 14*/*20, whereas the biology-oriented Overall(Bio) rises in 10*/*20 and falls in 10*/*20.

Latent dimensionality shows the clearest positive signal for batch removal and a modest benefit for the combined overall. Aggregating stepwise comparisons 10→20→30→40→50 at fixed (*n*_hidden_, *n*_layers_, Features), Overall increases in 27*/*48 steps, Overall(BC) in 35*/*48, and Overall(Bio) in 26*/*48. The endpoint contrast 50 −10 reinforces this: Overall improves in 8*/*12 matched triplets, Overall(BC) in 11*/*12, and Overall(Bio) in 6*/*12. Increasing *n*_latent_ therefore clearly helps batch correction and typically nudges the primary Overall upward; the impact on biology is mixed-to-slightly-positive on average.

Feature selection favors Full on this dataset, especially for batch. In paired HVG−Full comparisons at fixed (*n*_hidden_, *n*_latent_, *n*_layers_), Overall is higher with HVG in 11*/*30 and lower in 19*/*30 pairs, Overall(BC) is higher in 2*/*30 and lower in 28*/*30, and Overall(Bio) is higher in 10*/*30 and lower in 20*/*30. Thus, Full generally outperforms HVG for the composite scores on Zenodo 11100300, particularly for batch-oriented performance. However, the single best Overall configuration in this grid is HVG: 0.78615 at (256, 30, 1, HVG). In practice, Full is the stronger default across matched settings, but careful tuning (e.g., shallow depth with *n*_latent_ ≈30) can yield an HVG peak.

#### Per-Metric Performance on ZENODO 11100300 (18 PBMC SAMPLES) - MrVI

Table 9 reports the MrVI sweep on Zenodo 11100300 over *n*_hidden_∈ {128, 256}, *n*_latent_ ∈{10, 20, 30, 40, 50}, *n*_layers_ ∈{1, 2, 3}, and Features ∈{Full, HVG}. The best Overall is 0.75718 at (256, 50, 1, Full), while the worst Overall is 0.70797 at (256, 20, 3, Full).

**Table 9.**
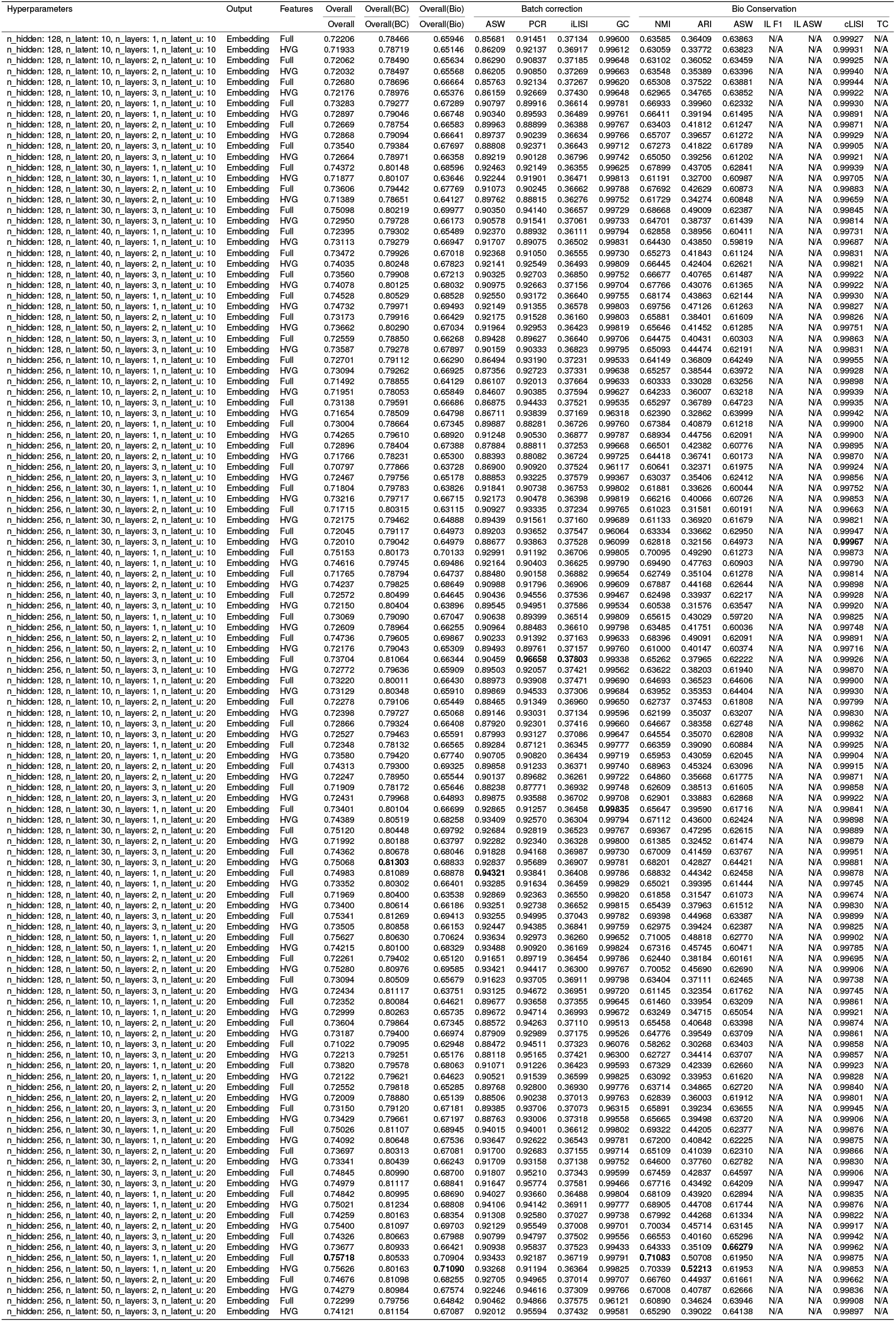
Performance of MrVI on the Zenodo 11100300 (18 PBMC Samples)

As shown in the cross-metric bar chart provided in Figure 23, changing hidden size from 128 to 256 yields small but consistent gains across the composites. In matched pairs that fix (*n*_latent_, *n*_layers_, Features), Overall improves in 17*/*30 and declines in 13*/*30 pairs, Overall(BC) improves in 19*/*30 and declines in 11*/*30, and Overall(Bio) improves in 17*/*30 and declines in 13*/*30. Thus, larger hidden size modestly lifts both batch and biology composites and nudges the primary Overall upward.

Increasing depth from *n*_layers_=1 to 3 is broadly unfavorable for the aggregate scores. At fixed (*n*_hidden_, *n*_latent_, Features), Overall rises in 7*/*20 and falls in 13*/*20 pairs, Overall(BC) is essentially flat with 10*/*20 up and 10*/*20 down, and Overall(Bio) rises in 7*/*20 and falls in 13*/*20. Deeper encoders, therefore, tend to reduce biological conservation and the primary Overall, while not improving the batch composite on average.

Latent dimensionality shows a clear preference toward higher capacity in this grid. Although the table does not contain enough perfectly matched stepwise chains to report full 10→ 20 →30→ 40→ 50 counts at fixed (*n*_hidden_, *n*_layers_, Features), the strongest configurations concentrate at larger *n*_latent_: the best Overall occurs at *n*_latent_=50 with shallow depth, and several batch- and clustering-oriented peaks are also at higher latent values.

Stepping *n*_latent_u_ from 10 to 20 yields consistent gains on this dataset. In paired comparisons at fixed (*n*_hidden_, *n*_latent_, *n*_layers_, Features), the primary Overall increases in 40*/*60 pairs, the batch composite Overall(BC) in 51*/*60, and the biology composite Overall(Bio) in 32*/*60. Batch metrics benefit strongly: Batch ASW rises in 54*/*60 and PCR in 50*/*60; iLISI shifts only slightly on average (up 24*/*60). Clustering agreement improves as well: NMI up 31*/*60 and ARI up 34*/*60. Label compactness also trends upward (Label ASW up 42*/*60), while graph connectivity is nearly unchanged on average (up 32*/*60, 1 flat) and cLISI is essentially flat (up 29*/*60).

By feature set, the *u* increase helps both Full and HVG, with a stronger batch gain under HVG. For Full: Overall up 18*/*30, Overall(BC) up 23*/*30, Overall(Bio) up 15*/*30. For HVG: Overall up 22*/*30, Overall(BC) up 28*/*30, Overall(Bio) up 17*/*30. In short, enlarging *n*_latent_u_ from 10 to 20 reliably improves batch removal and usually lifts the primary Overall, while also providing modest gains in clustering agreement and biology; neighborhood-level batch mixing (iLISI) moves only slightly on average.

Metric-wise optima align with these trends. Overall peaks at (256, 50, 1, Full) with 0.75718; the batch composite Overall(BC) peaks at (128, 30, 3, HVG) with 0.81304; the biology composite Overall(Bio) peaks at (256, 50, 1, HVG) with 0.71090. Among individual metrics: Batch ASW is highest at (128, 40, 1, Full) with 0.94321; PCR and iLISI peak at (256, 50, 3, Full) with 0.96658 and 0.37803, respectively; graph connectivity is highest at (128, 30, 1, Full) with 0.99835; clustering agreement is strongest at high latent with shallow depth, with NMI peaking at (256, 50, 1, Full) with 0.71083 and ARI at (256, 50, 1, HVG) with 0.52213; Label ASW peaks at (256, 40, 3, HVG) with 0.66279; cLISI peaks at (256, 30, 3, HVG) with 0.99967.

#### Per-Metric Performance on ZENODO 11100300 (18 PBMC SAMPLES) - LDVAE

Table 10 reports the LDVAE sweep on Zenodo 11100300 over *n*_hidden_ ∈{128, 256}, *n*_latent_∈{10, 20, 30, 40, 50}, *n*_layers_ ∈{1, 2, 3}, and Features ∈{Full, HVG}. The best Overall is 0.74219 at (256, 50, 1, HVG), while the worst Overall is 0.60572 at (256, 20, 3, Full).

**Table 10.**
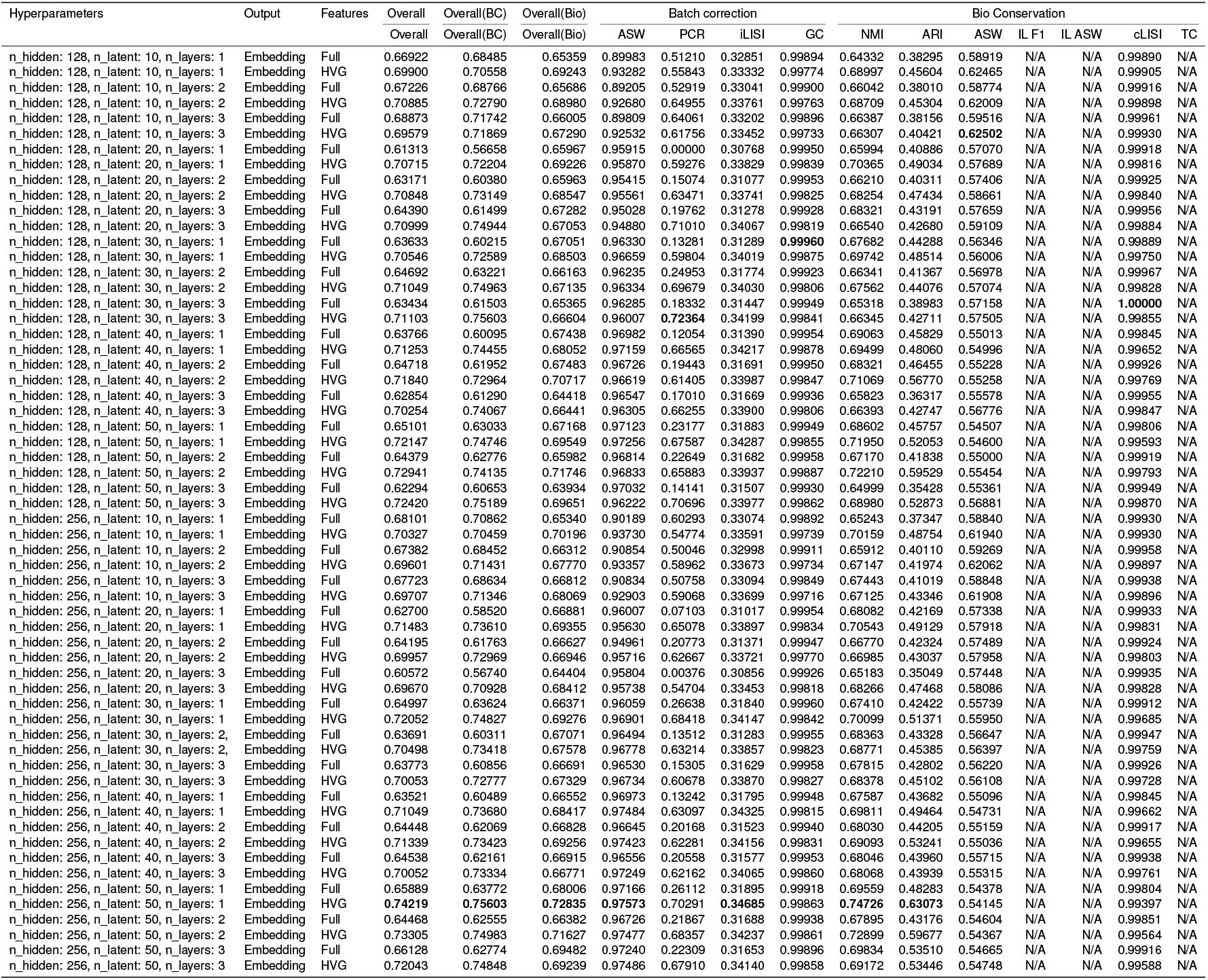
Performance of LDVAE on the Zenodo 11100300 (18 PBMC Samples)

As shown in the cross-metric bar chart provided in Figure 24, changing hidden size from 128 to 256 yields a small net gain in the primary aggregate and a clearer lift in biology, with a slight reduction in the batch composite. In paired comparisons that fix (*n*_latent_, *n*_layers_, Features), Overall rises in 16*/*30 and falls in 14*/*30 pairs, Overall(BC) rises in 14*/*30 and falls in 16*/*30, and Overall(Bio) rises in 20*/*30 and falls in 10*/*30.

Depth is broadly unfavorable for the main and biology composites. From *n*_layers_=2 versus 1, Overall is up 8*/*20 and down 12*/*20, Overall(BC) is essentially flat with 10*/*20 up, and Overall(Bio) is up 5*/*20 and down 15*/*20. From *n*_layers_=3 versus 1, Overall is up 7*/*20 and down 13*/*20, Overall(BC) is up 11*/*20, and Overall(Bio) is up 1*/*20 and down 19*/*20. Deeper encoders, therefore, tend to reduce biological conservation and slightly depress the primary. Overall, the batch composite is flat to modestly positive only at three layers.

Latent dimensionality shows the clearest positive signal for batch removal and a modest lift for the primary aggregate. Stepwise across 10→ 20 →30 →40 →50 at fixed (*n*_hidden_, *n*_layers_, Features), Overall increases in 31*/*48 steps, Overall(BC) in 40*/*48, while Overall(Bio) increases in 18*/*48. The endpoint contrast 50− 10 reinforces this: Overall improves in 8*/*12 matched triplets, Overall(BC) in 10*/*12, and Overall(Bio) in 6*/*12. Increasing *n*_latent_ thus reliably helps batch correction and typically nudges the primary Overall upward; the biology composite often softens stepwise but is roughly balanced end-to-end.

Feature selection favors HVG on this dataset. In paired HVG− Full comparisons at fixed (*n*_hidden_, *n*_latent_, *n*_layers_), Overall is higher with HVG in 16*/*30 and lower in 14*/*30, Overall(BC) is higher in 16*/*30, and Overall(Bio) is higher in 18*/*30. HVG is therefore modestly better for the primary, batch, and biology composites in this grid.

Metric-wise optima are consistent with these trends. Overall peaks at (256, 50, 1, HVG) with 0.74219; the batch composite Overall(BC) peaks at (128, 30, 3, HVG) with 0.75603; the biology composite Overall(Bio) peaks at (256, 50, 1, HVG) with 0.72835. Among individual metrics: Batch ASW is highest at (256, 50, 1, HVG) with 0.97573; PCR peaks at (128, 30, 3, HVG) with 0.72364; iL-ISI peaks at (256, 50, 1, HVG) with 0.34685; graph connectivity is highest at (128, 30, 1, Full) with 0.99960; clustering agreement is strongest at high latent with shallow depth, with NMI and ARI both peaking at (256, 50, 1, HVG) with 0.74726 and 0.63073, respectively; Label ASW peaks at (128, 10, 3, HVG) with 0.62502; cLISI peaks at (128, 30, 3, Full) with 1.00000.

## Computational Complexity

Each fully connected (dense) layer is implemented as a general matrix–matrix multiplication (GEMM). For a mini-batch of size *B*, input width *in*, and output width *out*, the forward/backward work scales as𝒪 (*B*.*in*.*out*)[56]. Consequently, a layer that maps latent *z* to hidden *H* (or vice versa) with *B* samples costs 𝒪 (*BHz*). Stacking hidden layers adds additional GEMMs: an *H*→ *H* block contributes 𝒪 (*BH*^2^) per layer, while gene-facing heads *H* → *G* cost 𝒪 (*BHG*). Hence, increasing *n*_latent_ affects only the *H* ↔ *z* interfaces (linear in *z*), whereas the dominant terms are the gene heads *H* → *G* and any extra hidden layers *H* → *H*.

### Practical compute implications of tuning *z, H, G*, and *B*

Each fully connected (dense) layer is implemented as a general matrix–matrix multiplication (GEMM). For a mini-batch of size *B*, input width *in*, and output width *out*, the forward/backward work scales as 𝒪 (*B*.*in*.*out*)[56]. We use this rule to reason about how training cost changes when we adjust the latent size *z*, hidden width *H*, the number of genes *G*, and the batch size *B*. Throughout, we keep in mind the typical magnitudes in single-cell integration: *G* in the *thousands, H* in the *hundreds*, and *z* in the *tens*. In practice, both *H* and *G* tend to move in large steps (e.g., *H* : 128→ 256; *G*: all-genes vs. ∼5,000 HVGs), and *B* is usually chosen to saturate memory, so the most nimble knob is *z*.

### scVI (MLP decoder, *z* → *H* → *G*)

With at least one hidden layer in the decoder, the terms that depend on *z* are confined to the *H*↔*z* interfaces: the encoder heads *H* →*z* and the decoder entrance *z*→*H*, giving an incremental cost.

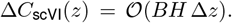

The dominant gene-facing multiplies *G*→*H* (encoder front) and *H*→*G* (decoder head) remain𝒪 (*BHG*) and do not grow with *z*. Hence the fractional overhead of raising *z* by Δ*z* is approximately

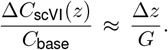

Since *G* is in the thousands and Δ*z* is in the tens, this overhead is sub-percent in typical settings. Conclusion: for scVI, increasing *z* is the cheapest way to add capacity and increase performance as proved by our extensive experiments; it barely perturbs the big𝒪 (*BHG*) terms. This makes scVI particularly attractive when one wants to improve integration quality with minimal training-time impact.

### LDVAE and MrVI (linear gene projection; decoder cost includes *BGz*)

LDVAE replaces the non-linear decoder with a single linear map *z*→*G*, and MrVI’s generative path also produces *G*-dimensional outputs via a linear projection of a *z*-width representation. In both cases, the decoder contributes a term 𝒪 (*BGz*). The encoder front *G* →*H* remains𝒪(*BGH*). Consequently, the leading-order training cost can be summarized as

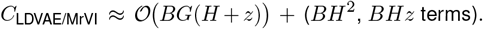

A small increase in *z* yields

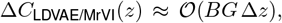

while a small increase in *H* yields

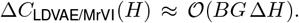

*Per unit of width*, Δ*z* and Δ*H* cost the same order: both scale like *BG*. What makes *z* cheaper *in practice* is the size of the step: we usually take Δ*z* in the tens, whereas Δ*H* is commonly a large jump (e.g., +128). Formally, the fractional overheads satisfy

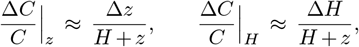

so for typical magnitudes (*H* hundreds, *z* tens) a +10–+40 change in *z* is usually less expensive than a +128 change in *H*, even though *z* appears in a “big” *BGz* term. By contrast, changing *G* or *B* directly scales every large multiply (∝*BG*), so those knobs are inherently costly; therefore, restricting to HVGs (smaller *G*) with tuning *z* is such a powerful lever for LDVAE/MrVI.

### Memory footprint

In scVI, raising *z* only widens the *H* ↔*z* blocks, adding𝒪 (*H* Δ*z*) parameters. In LDVAE/MrVI, the gene projection widens with *z*, adding 𝒪 (*G* Δ*z*) parameters, which are manageable with HVGs, but much larger if using all genes.

### Guidance synthesizing magnitudes and step sizes

Because *H* and *G* typically move in large chunks and *B* multiplies all costs, the small and targeted adjustments available through *z* make it the most economical capacity knob across models for different reasons: in scVI it is intrinsically cheap (Δ*T* ∝*BH* Δ*z* while 𝒪 (*BHG*) is unchanged), and in LDVAE/MrVI it is cheaper in practice because Δ*z* (tens) is much smaller than common Δ*H* (hundreds) or changes to *G* (thousands). Accordingly, our practical recipe is: use HVGs to control *G*, keep networks shallow, and treat *z* as the first tuning knob. In particular, scVI remains the go-to model when one wants strong integration with minimal incremental training cost from increasing latent dimensionality.

### Score-Complexity Trade-offs

Figure 11 summarizes the effect of varying a single hyper-parameter while holding the others fixed. The first panel reports the mean score, and the second reports the mean training time, both averaged across datasets. Across models, increasing the Z-latent dimensionality *n*_latent_*Z* yields the most favorable score–cost trade-off, especially for scVI, typically improving performance with the smallest increase in memory and only modest growth in time. Pairwise hyperparameter comparisons further show in detail that raising *n*_latent_*Z* benefits most individual metrics and outperforms alternatives such as increasing depth or hidden width in terms of score–time/memory efficiency.

**Figure 11.**
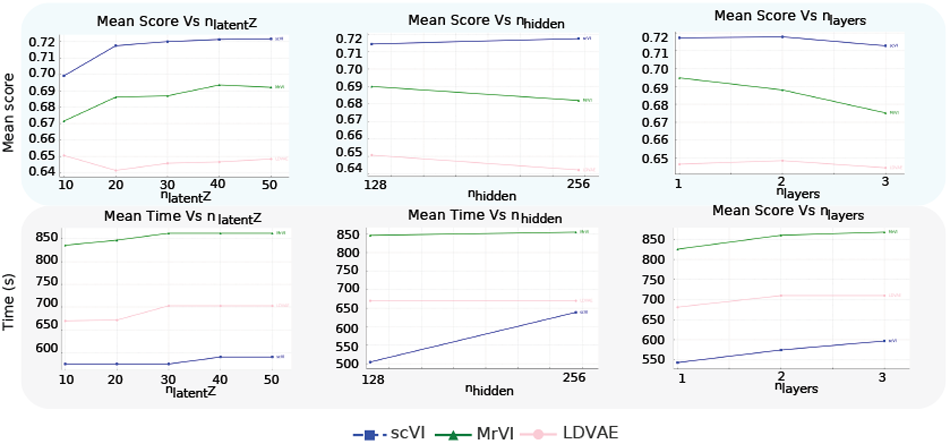
Cross-dataset trends in integration performance and runtime across architectural hyperparameters. Mean overall integration score (y axis; blue panels) as a function of shared latent dimensionality (nlatent,Z; left), hidden-layer width (nhidden; middle) and network depth (nlayers; right). Mean training time per run (y axis; grey panels) as a function of nlatentZ, nhidden and nlayers. In each plot, one architectural hyperparameter is varied while the remaining two are held fixed (as in the benchmark grid; Table 1), and values are averaged across datasets. Lines correspond to model families (scVI, MrVI and LDVAE), summarising how performance and computational cost change as model capacity increases.

## Supplementary Note 1: Data Availability and Reproducibility

Benchmarking was done with Snakemake v9.11.5 to ensure a reproducible and scalable analysis pipeline. the complete workflow including environment and code is available in https://github.com/Kassab11/A-hyperparameter-benchmark-of-VAE-based-methods-for-scRNA-seq-batch-integration. We also provide an interactive Jupyter notebook for hyperparameter sweeps in which users can vary *n*_latent_(*z*), *n*_latent_(*u*), *n*_hidden_, *n*_layers_, and the feature set; plots update in real time to streamline comparison and guide selection of configurations for their study.

We conducted all experiments on Ubuntu 22.04 LTS with an NVIDIA RTX 4090 (24 GB VRAM) and 128 GB RAM, Python v3.12.2 (https://www.python.org); Scanpy v1.11.0 for preprocessing and visualization (https://scanpy.readthedocs.io); scvi-tools v1.3.0 for probabilistic modeling with scVI, MrVI, and LDVAE (https://docs.scvi-tools.org); PyTorch v2.6.0+cu124 as a primary deep-learning backend (https://pytorch.org); JAX v0.4.35 (Google AI, https://github.com/google/jax); scIB v1.1.7 for integration and conservation metrics (https://scib.readthedocs.io), and snakemake v9.11.5 (https://snakemake.readthedocs.io/en/stable/).

This study uses four single-cell RNA sequencing datasets that span various experimental conditions and batch structures. The first dataset is the Human Immune Cells dataset from the Open Problems in Single-Cell Analysis initiative [1], accessible at https://openproblems.bio/datasets/openproblems_v1/immune_cells. It includes 33,506 immune cells and 12,303 genes collected from peripheral blood and bone marrow, distributed across ten batches and generated using both 10X Genomics and Smart-seq2 technologies. In addition, two datasets were obtained from Zenodo upon request. The 24 PBMC dataset contains 76,535 cells and 36,601 genes collected from capillary blood samples across 14 batches using 10X Genomics [57], accessible upon request at https://zenodo.org/records/8020792. The 18 PBMC dataset includes 55,260 cells and 36,601 genes from four batches, corresponding to samples collected during a remission-inducing antibiotic intervention [58] accessible upon request at https://zenodo.org/records/11100300. The Tabula Muris atlas is a cross-tissue mouse single-cell RNA-seq resource profiling on the order of 10^5^ cells across roughly 20 organs using two complementary modalities, droplet-based 3^’^ UMI (10x) for breadth and FACS/Smart-seq2 [51]. The dataset includes 356,213 cells and 20,116 genes. The atlas includes 51 immune cell-type labels (e.g., T/B/NK, myeloid and dendritic subsets, tissue-resident macrophages) and 103 non-immune labels (epithelial, endothelial/vascular, stromal/mesenchymal, neural, muscle, hepatic, renal, pancreatic, etc.), for a total of about 154 distinct labels. The dataset can be obtained from https://biohub.org/sf/tabula-muris/

## Supplementary Note 2: Dataset Statistics - Human Immune Dataset

**Figure 12.**
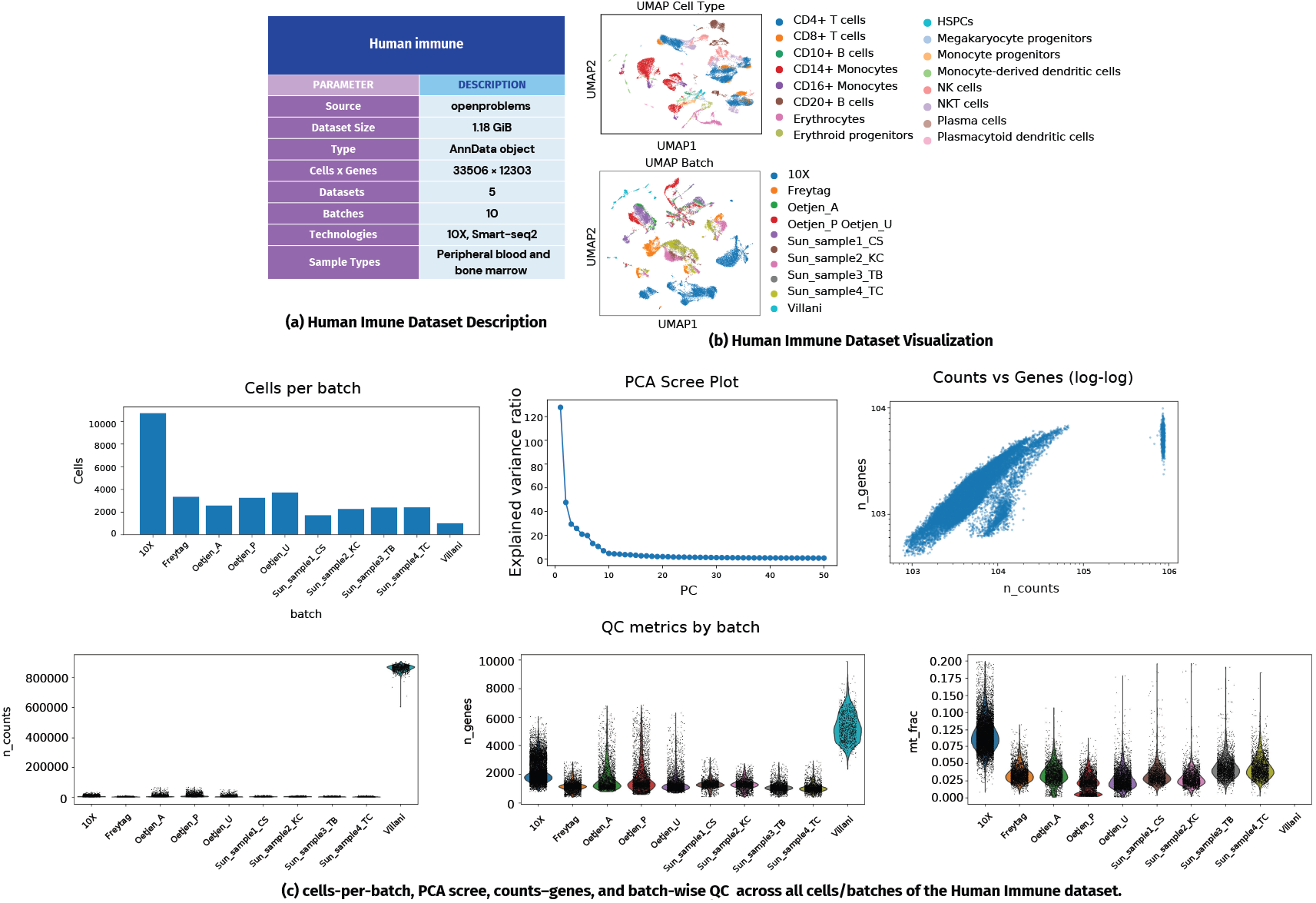
Summary of dataset description and visualization used in Human Immune Dataset. (a) OpenProblems immune dataset: 33,506 cells and 12,303 genes across 5 datasets and 10 batches, generated using 10X and Smart-seq2 technologies from peripheral blood and bone marrow samples. (b) UMAP projections of the OpenProblems dataset colored by cell type (top) and batch (bottom), highlighting biological diversity and batch heterogeneity. (c) PCA scree plot, counts–genes, and mitochondrial fraction (top), dataset-level QC summary (bottom).

## Supplementary Note 3: Dataset Statistics - 18 PBMC Samples

**Figure 13.**
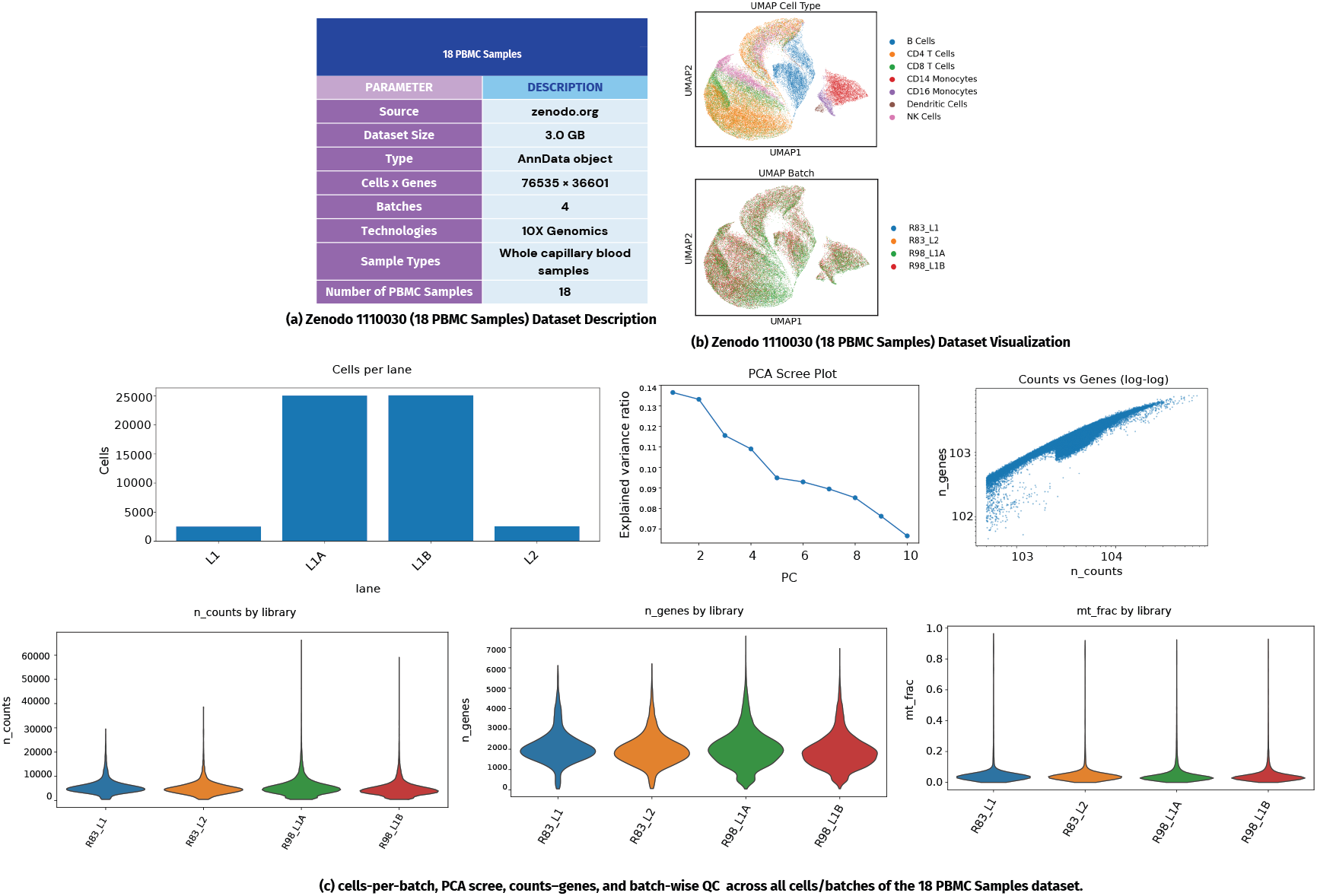
Summary of dataset description and visualization used in Zenodo 11100300 (18 PBMC Samples). (a) Zenodo 11100300 (18 PBMC Samples): 55,260 cells and 36,601 genes across 4 batches from whole capillary blood samples generated using 10X Genomics. (b) UMAP projections of Zenodo 8020792 (24 PBMC Samples) colored by cell type (top) and batch (bottom), showing pronounced batch effects and distinct cell type clustering. (c) PCA scree plot, counts–genes, and mitochondrial fraction (top), dataset-level QC summary (bottom).

## Supplementary Note 4: Dataset Statistics - 24 PBMC Samples

**Figure 14.**
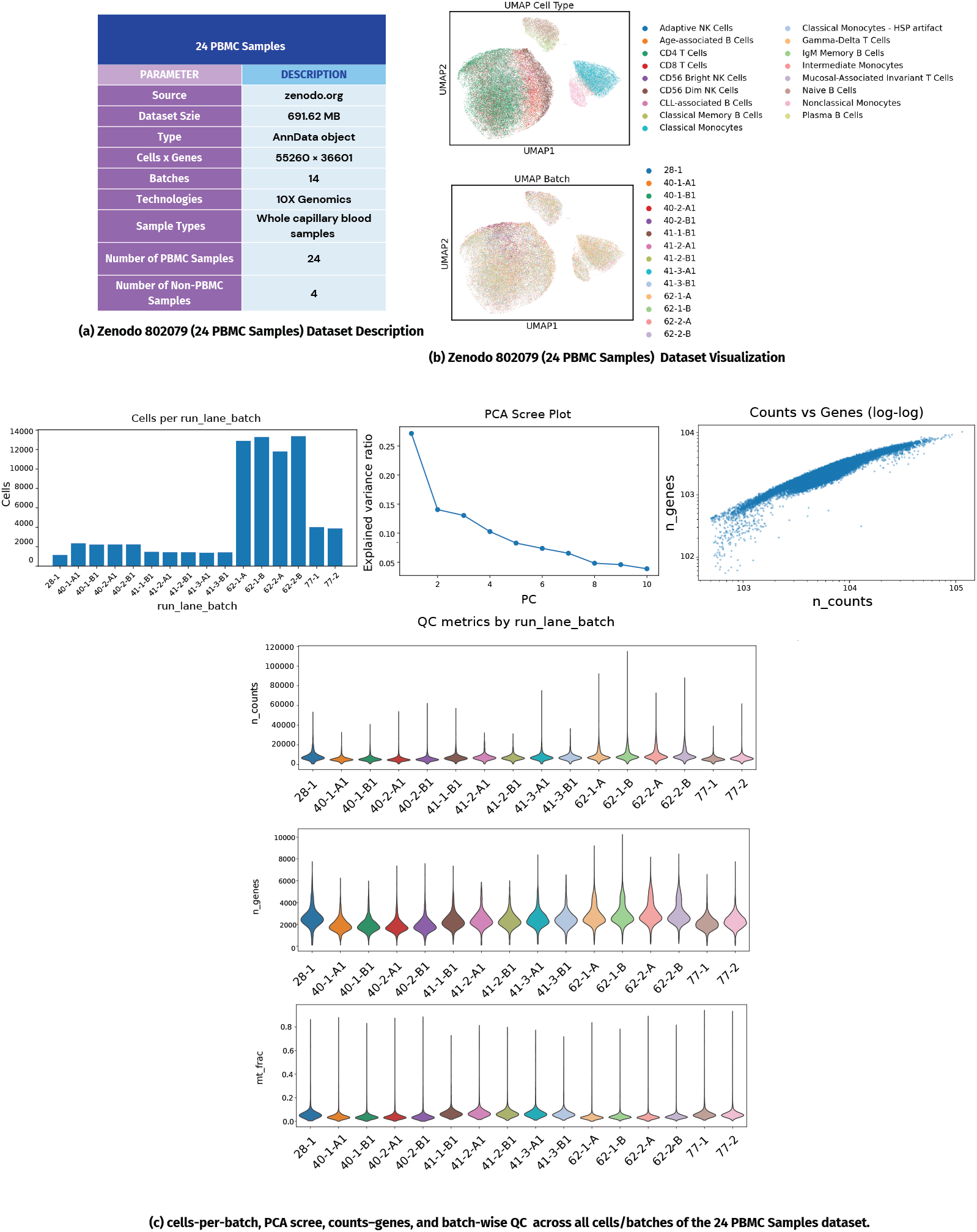
Summary of dataset description and visualization used in Zenodo 8020792 (24 PBMC Samples). (a) Zenodo 8020792 (24 PBMC Samples): 76,535 cells and 36,601 genes across 14 batches from whole capillary blood samples processed using 10X Genomics. (b) UMAP projections of Zenodo 8020792 (24 PBMC Samples) colored by cell type (top) and batch (bottom), showing pronounced batch effects and distinct cell type clustering. (c) PCA scree plot, counts–genes, and mitochondrial fraction (top), dataset-level QC summary (bottom).

## Supplementary Note 5: Dataset Statistics - Tabula Muris

**Figure 15.**
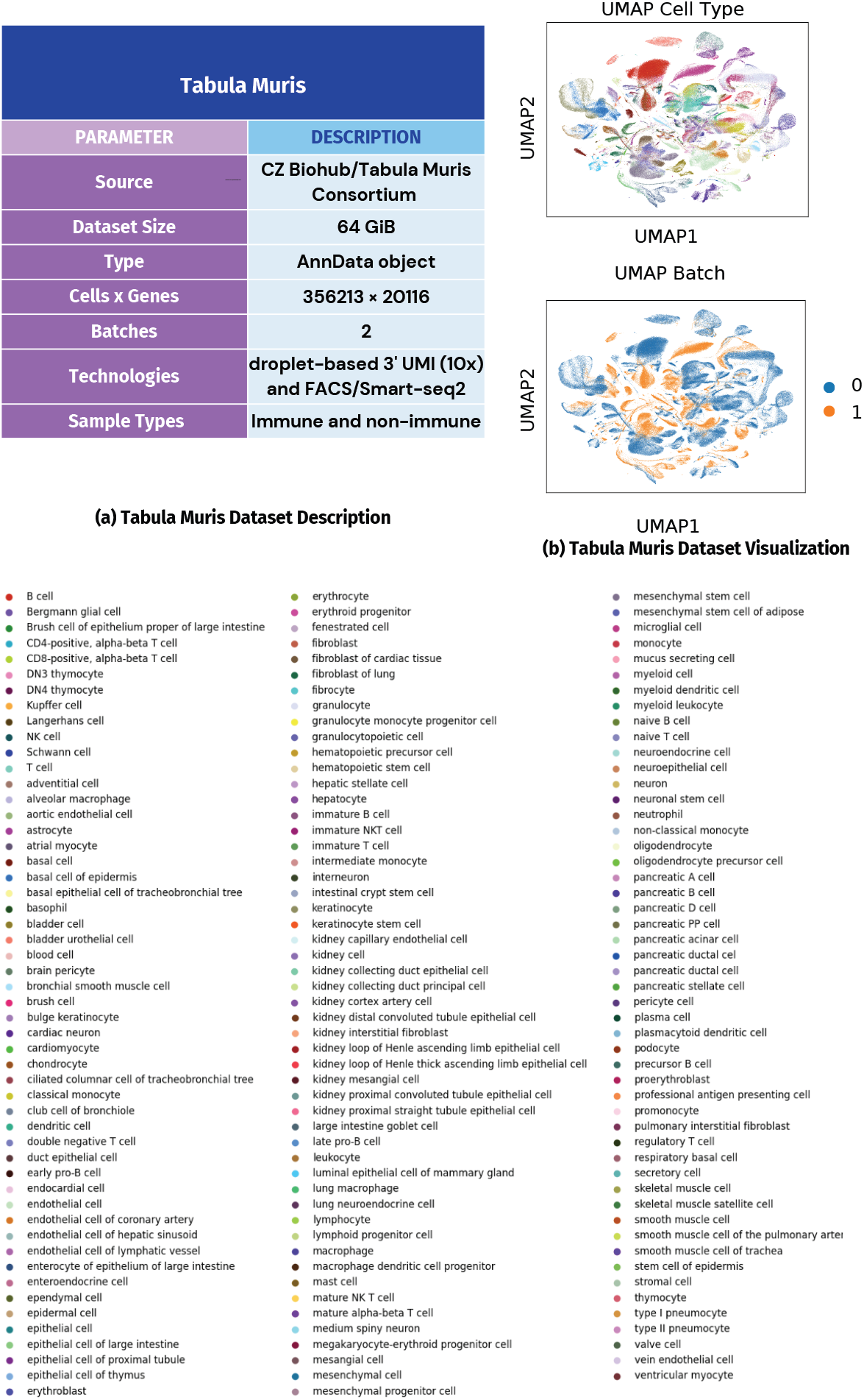
Summary of dataset description and visualization used in Tabula Muris. (a) Tabula Muris: 356,123 cells and 20,116 genes across 154 distinct cell types (b) UMAP projections of Tabula Muris colored by cell type (top) and batch (bottom).

## Supplementary Note 6: scVI Per Metric Performance on Human Immune

**Figure 16.**
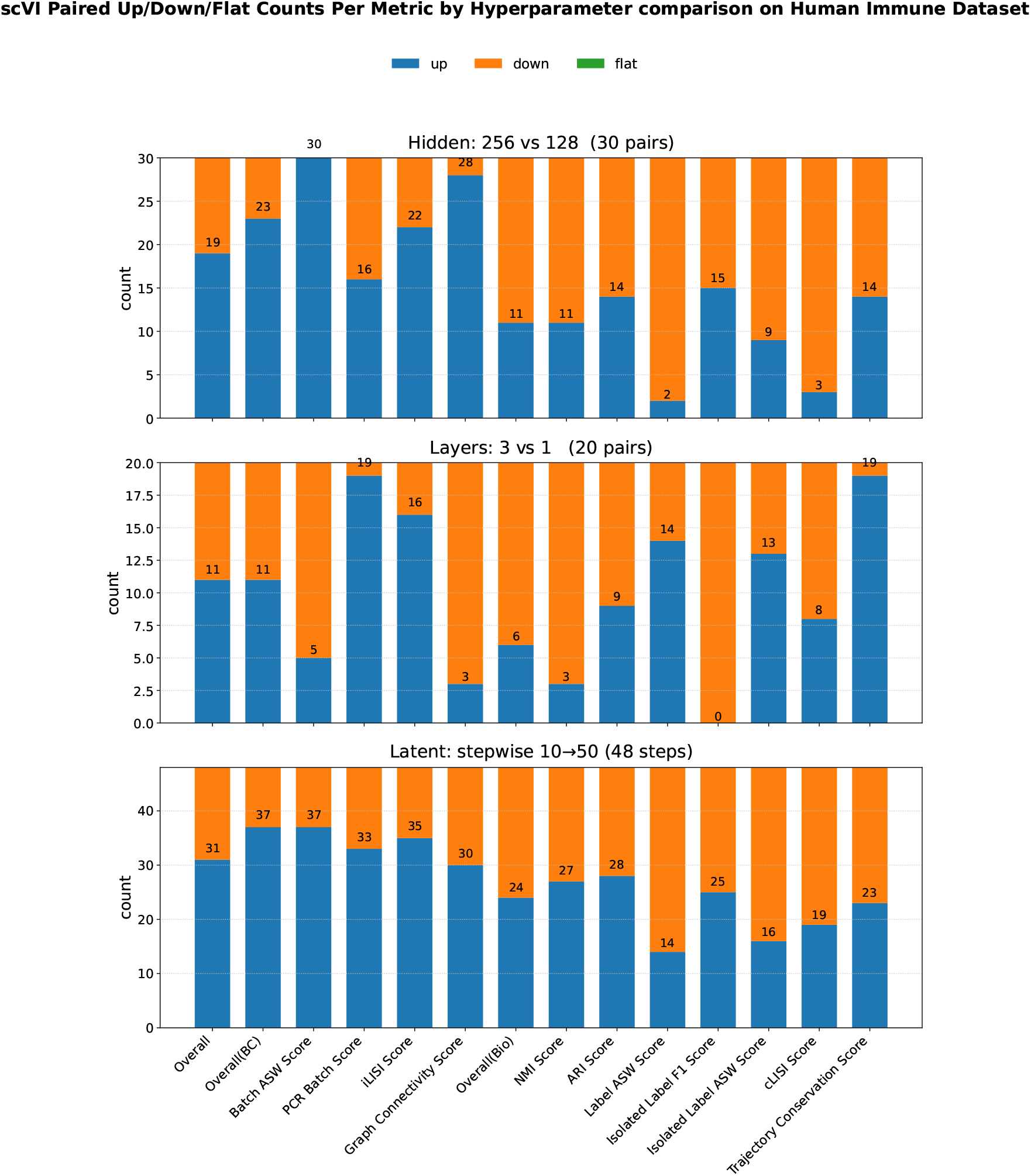
Paired comparison of scVI hyperparameters on the Human Immune dataset, showing the number of metrics that improve (“up”), decline (“down”), or remain unchanged (“flat”) when varying hidden units (256 vs. 128), network depth (3 vs. 1 layers), and latent dimensions (stepwise 10→50), evaluated on both full and HVG feature sets.

## Supplementary Note 7: MrVI Per Metric Performance on Human Immune

**Figure 17.**
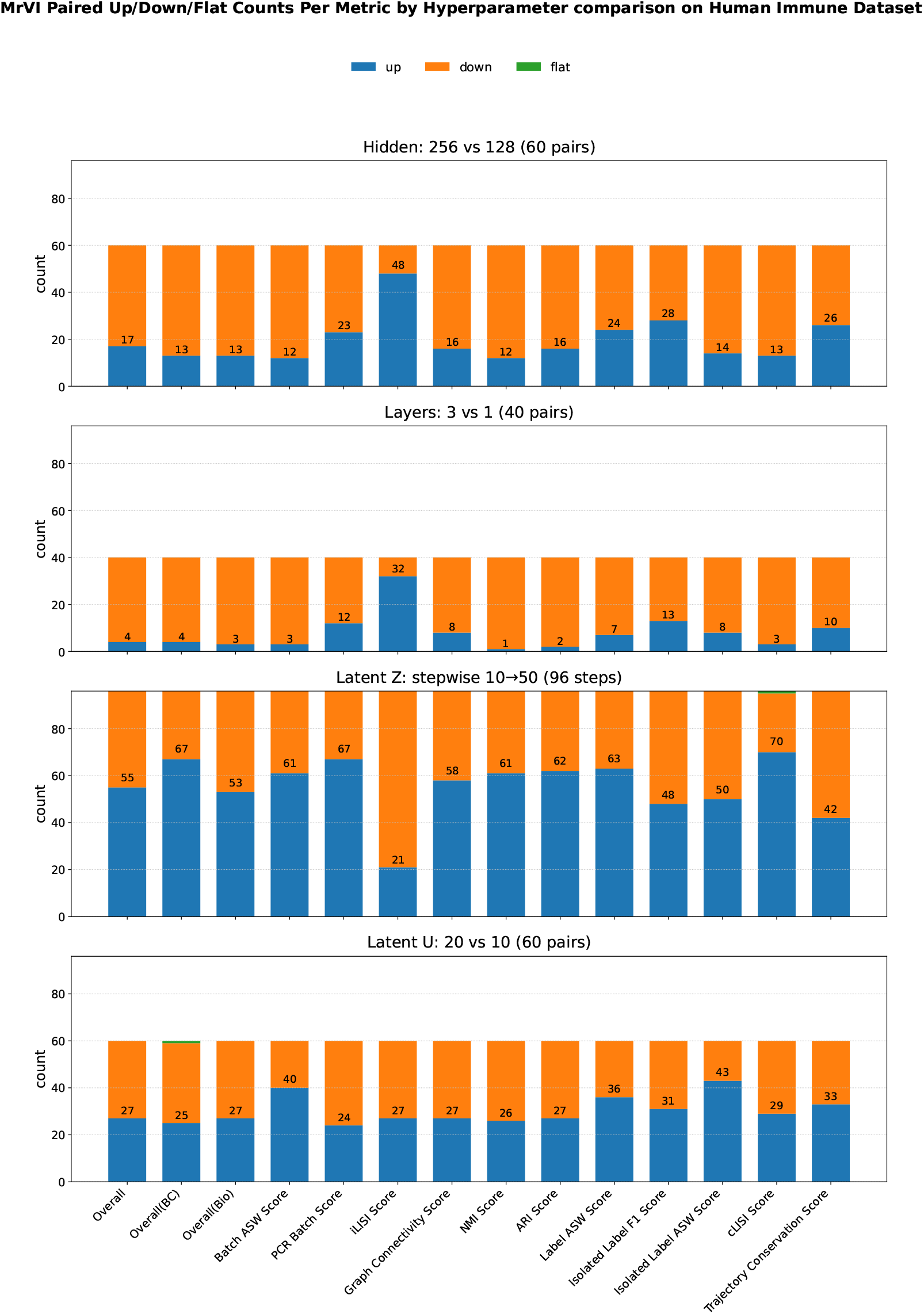
Paired comparison of MrVI hyperparameters on the Human Immune dataset, showing the number of metrics that improve (“up”), decline (“down”), or remain unchanged (“flat”) when varying hidden units (256 vs. 128), network depth (3 vs. 1 layers), latent dimension z (stepwise 10→50), and latent dimension u evaluated on both full and HVG feature sets.

## Supplementary Note 8: LDVAE Per Metric Performance on Human Immune

**Figure 18.**
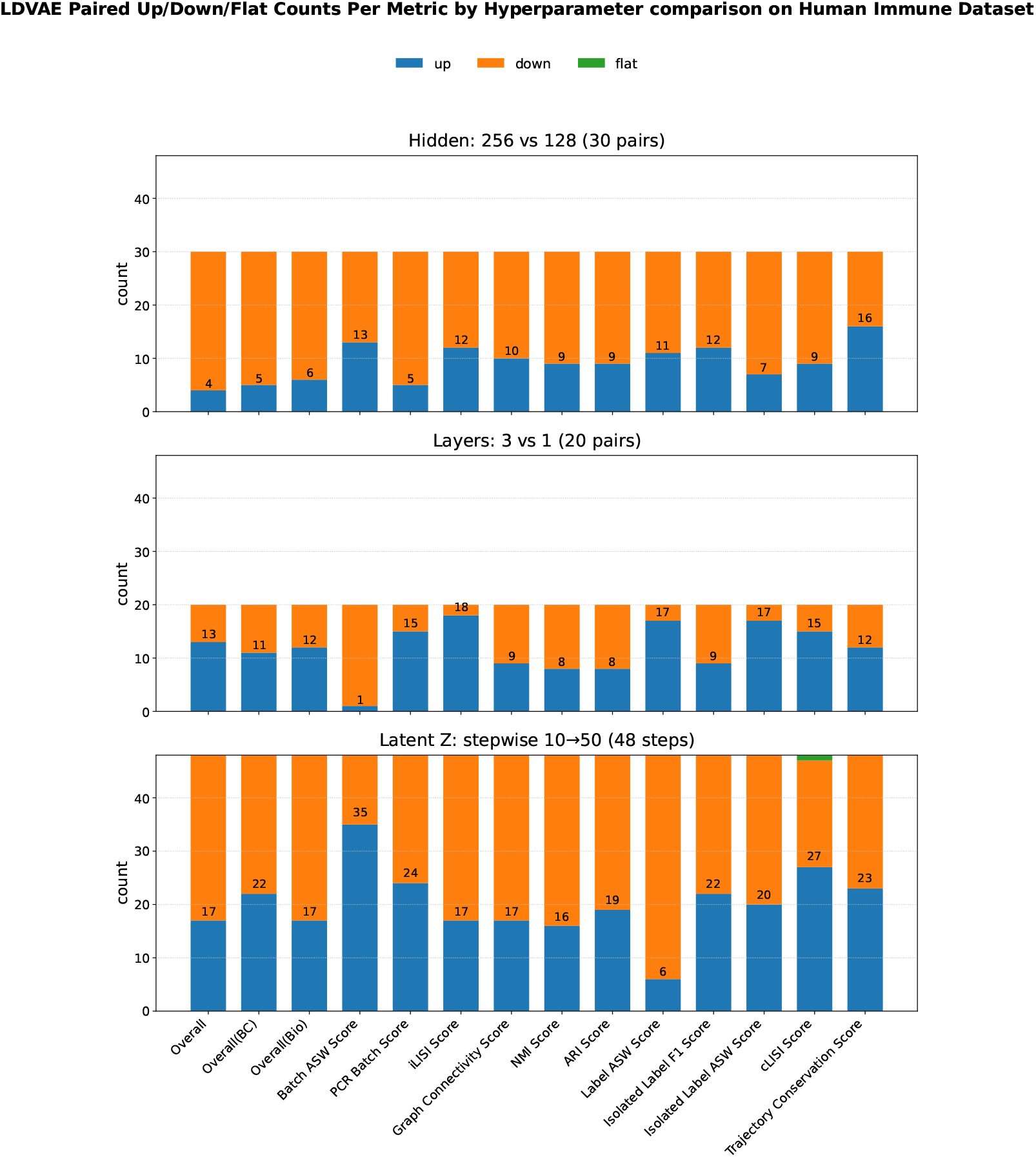
Paired comparison of LDVAE hyperparameters on the Human Immune dataset, showing the number of metrics that improve (“up”), decline (“down”), or remain unchanged (“flat”) when varying hidden units (256 vs. 128), network depth (3 vs. 1 layers), latent dimension z (stepwise 10→50), and latent dimension u evaluated on both full and HVG feature sets.

## Supplementary Note 9: scVI Per Metric Performance on Zenodo 8020792 (24 BMCSamples)

**Figure 19.**
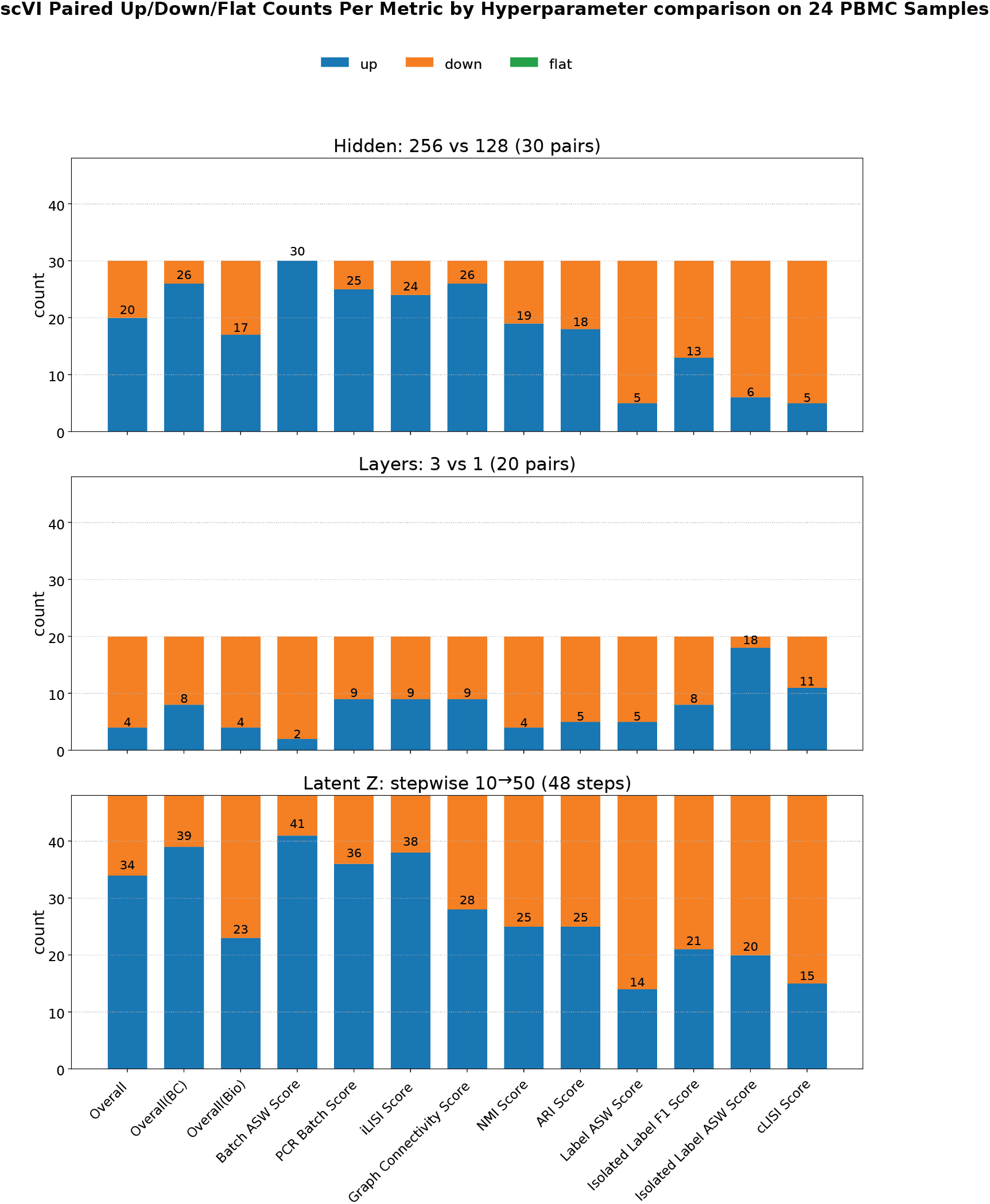
Paired comparison of scVI hyperparameters on the Zenodo 8020792 (24 PBMC Samples) dataset, showing the number of metrics that improve (“up”), decline (“down”), or remain unchanged (“flat”) when varying hidden units (256 vs. 128), network depth (3 vs. 1 layers), latent dimension z (stepwise 10→50), and latent dimension u evaluated on both full and HVG feature sets.

## Supplementary Note 10: MrVI Per Metric Performance on Zenodo 8020792 (24 PBMC Samples)

**Figure 20.**
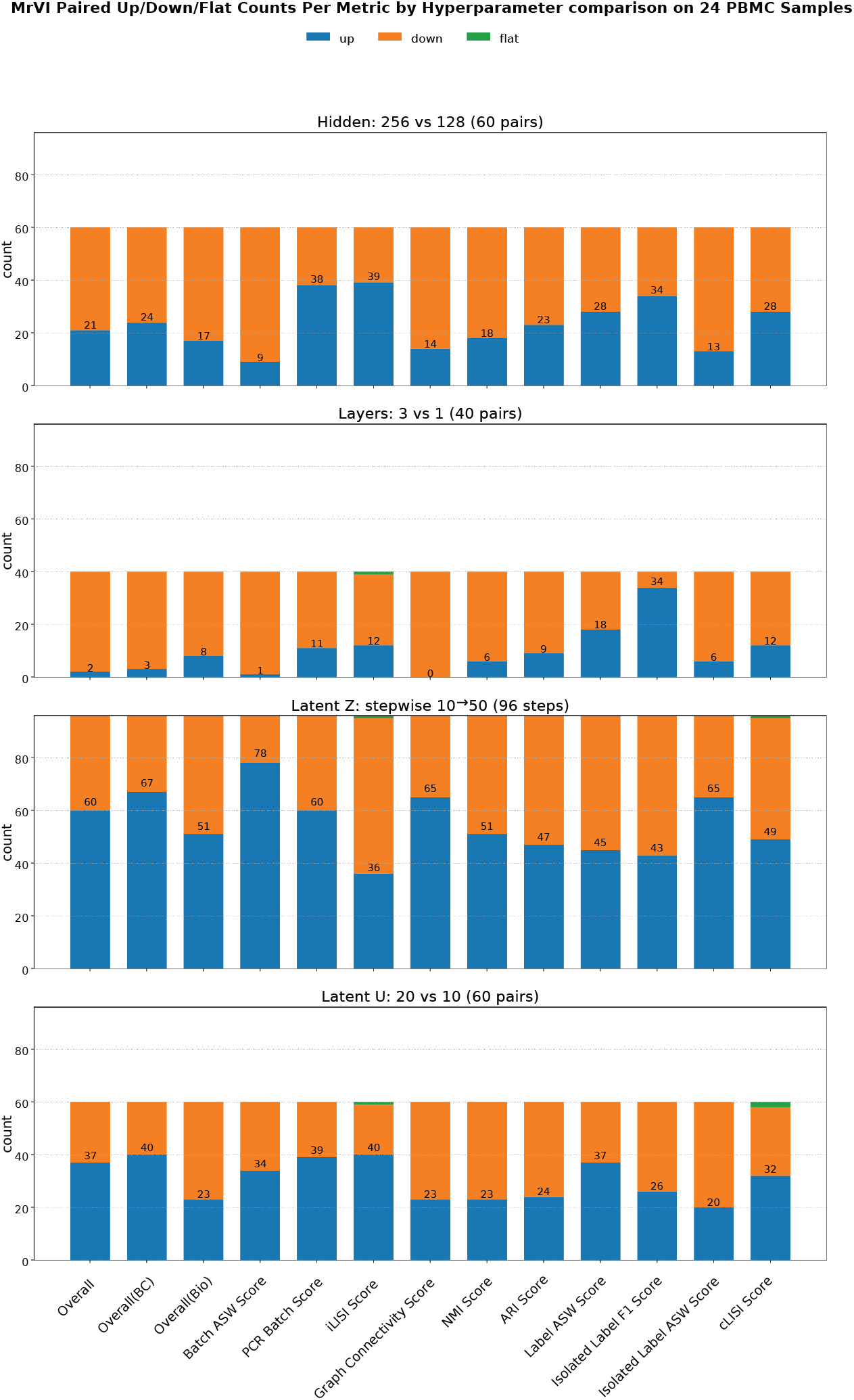
Paired comparison of MrVI hyperparameters on the Zenodo 8020792 (24 PBMC Samples) dataset, showing the number of metrics that improve (“up”), decline (“down”), or remain unchanged (“flat”) when varying hidden units (256 vs. 128), network depth (3 vs. 1 layers), latent dimension z (stepwise 10→50), and latent dimension u evaluated on both full and HVG feature sets.

## Supplementary Note 11: LDVAE Per Metric Performance on Zenodo 8020792 (24 PBMC Samples)

**Figure 21.**
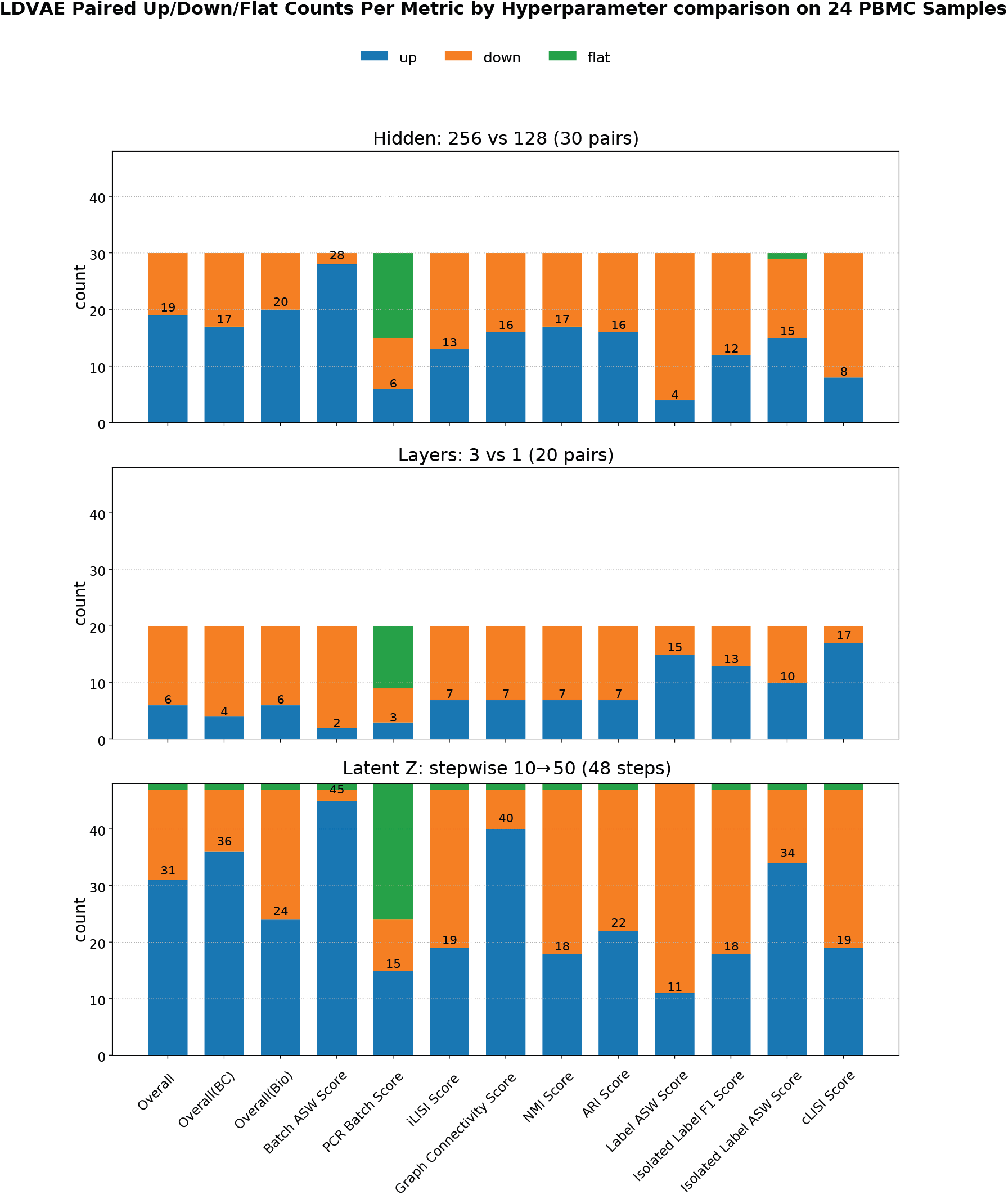
Paired comparison of LDVAE hyperparameters on the Zenodo 8020792 (24 PBMC Samples) dataset, showing the number of metrics that improve (“up”), decline (“down”), or remain unchanged (“flat”) when varying hidden units (256 vs. 128), network depth (3 vs. 1 layers), latent dimension z (stepwise 10→50) on both full and HVG feature sets.

## Supplementary Note 12: scVI Per Metric Performance on ZENODO 11100300 (18 PBMC SAMPLES)

**Figure 22.**
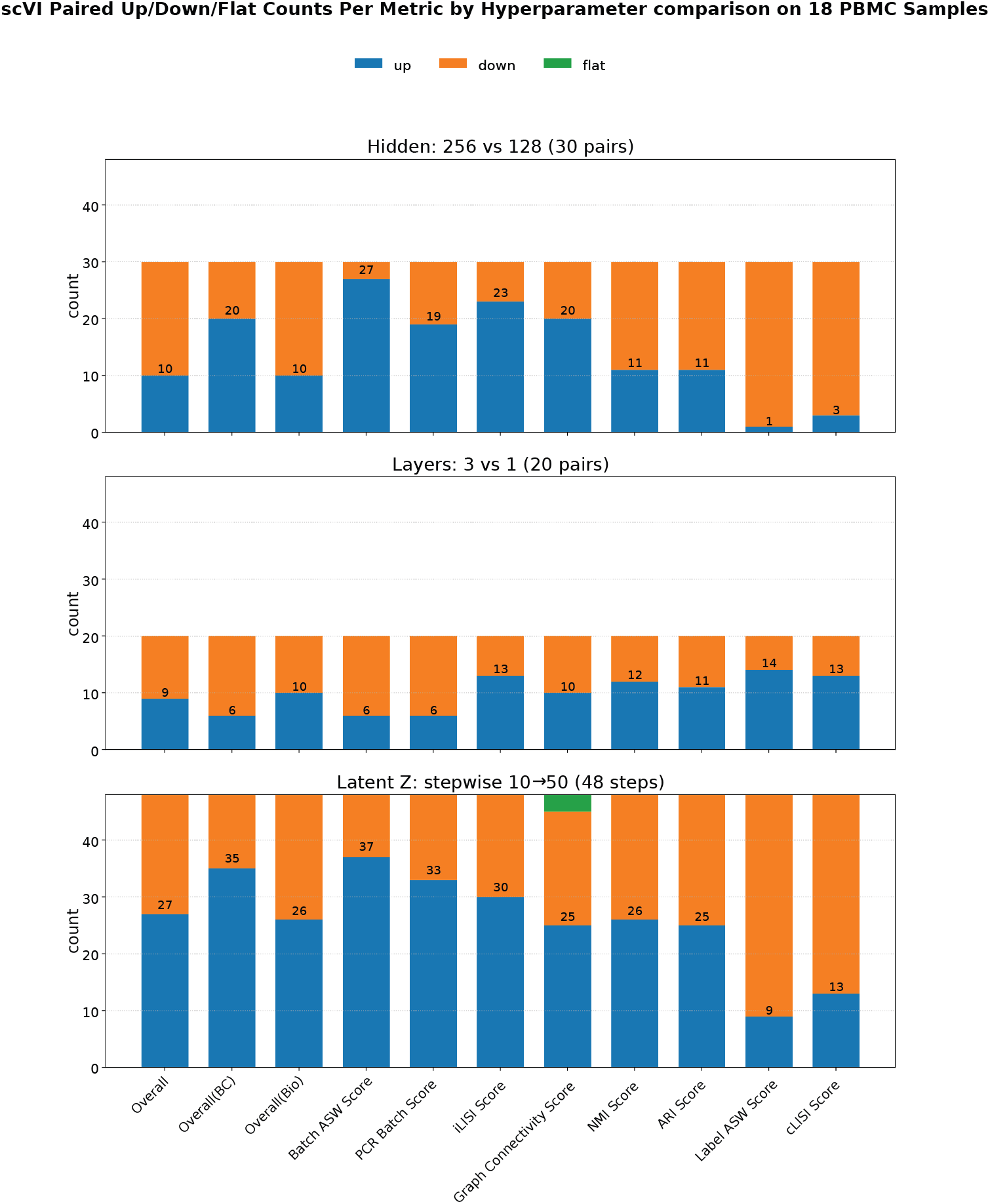
Paired comparison of scVI hyperparameters on the Zenodo 11100300 (18 PBMC Samples) dataset, showing the number of metrics that improve (“up”), decline (“down”), or remain unchanged (“flat”) when varying hidden units (256 vs. 128), network depth (3 vs. 1 layers), latent dimension z (stepwise 10→50) on both full and HVG feature sets.

## Supplementary Note 13: MrVI Per Metric Performance on ZENODO 11100300 (18 PBMC SAMPLES)

**Figure 23.**
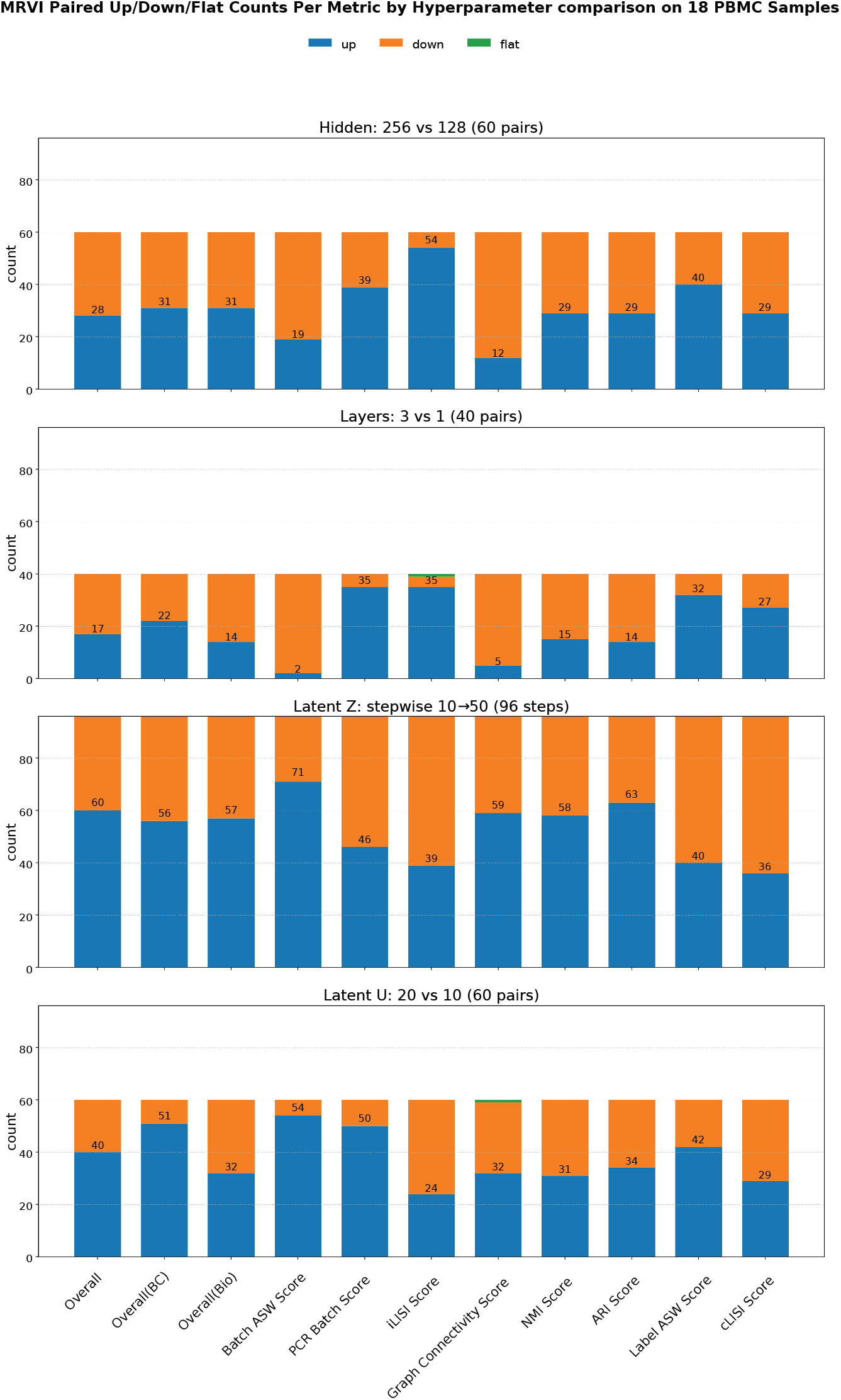
Paired comparison of MrVI hyperparameters on the Zenodo 11100300 (18 PBMC Samples) dataset, showing the number of metrics that improve (“up”), decline (“down”), or remain unchanged (“flat”) when varying hidden units (256 vs. 128), network depth (3 vs. 1 layers), latent dimension z (stepwise 10→50), and latent dimension u evaluated on both full and HVG feature sets.

## Supplementary Note 14: LDVAE Per Metric Performance on ZENODO 11100300 (18 PBMC SAMPLES)

**Figure 24.**
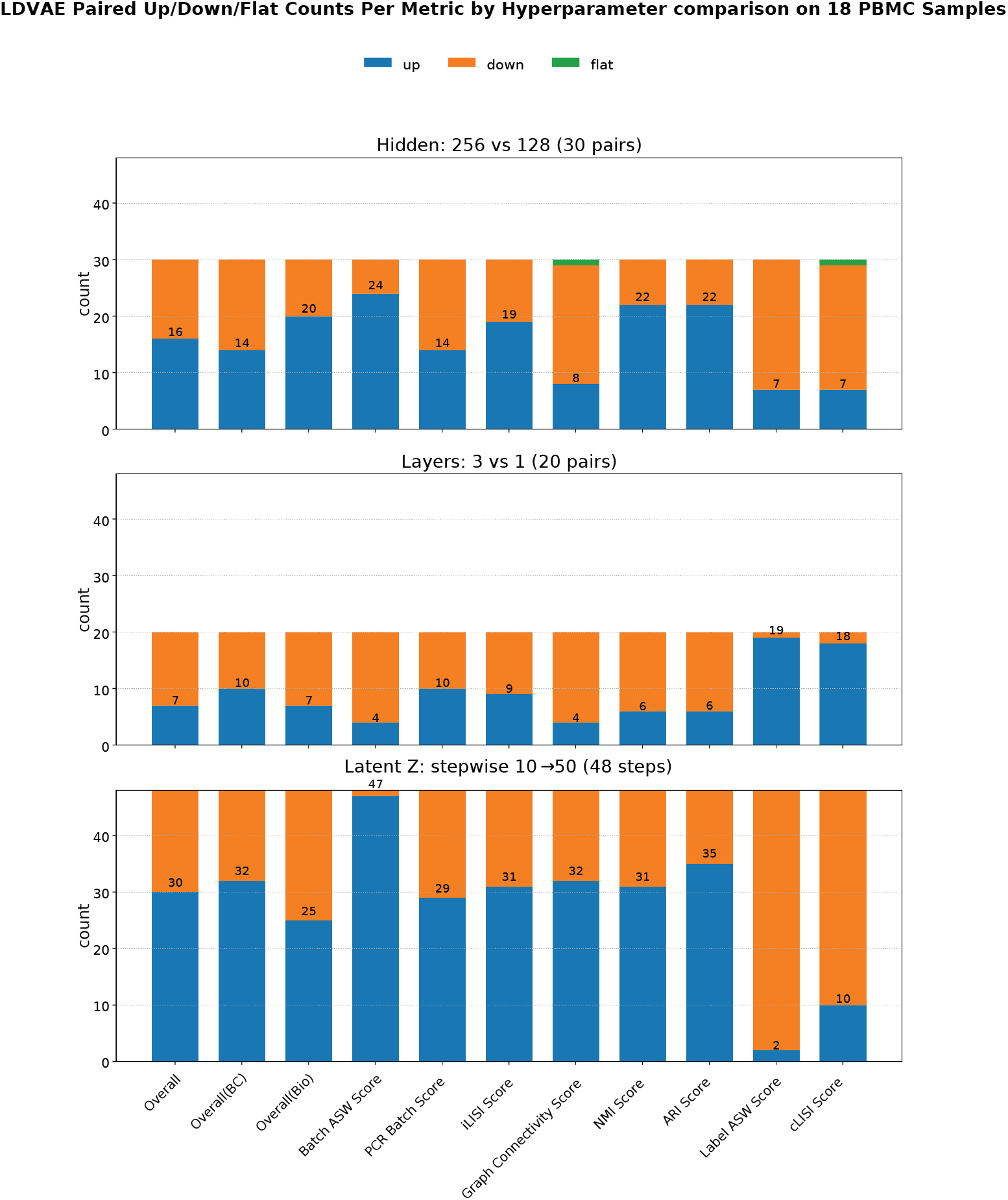
Paired comparison of LDVAE hyperparameters on the Zenodo 11100300 (18 PBMC Samples) dataset, showing the number of metrics that improve (“up”), decline (“down”), or remain unchanged (“flat”) when varying hidden units (256 vs. 128), network depth (3 vs. 1 layers), latent dimension z (stepwise 10→50) on both full and HVG feature sets.

